# Impact of prenatal delta-9-tetrahydrocannabinol exposure on mouse brain development: a fetal-to-adulthood magnetic resonance imaging study

**DOI:** 10.1101/2024.11.02.621669

**Authors:** Lani Cupo, Haley A. Vecchiarelli, Daniel Gallino, Jared VanderZwaag, Katerina Bradshaw, Annie Phan, Mohammadparsa Khakpour, Benneth Ben-Azu, Elisa Guma, Jérémie Fouquet, Shoshana Spring, Brian J. Nieman, Gabriel A. Devenyi, Marie-Eve Tremblay, M. Mallar Chakravarty

## Abstract

While cannabis use during pregnancy is often perceived as harmless, little is known about its consequences on offspring neurodevelopment. There is an urgent need to map the effects of prenatal cannabis exposure on the brain through the course of the lifespan. We used magnetic resonance imaging spanning nine timepoints, behavioral assays, and electron microscopy to build a trajectory from gestation to adulthood in mice exposed prenatally to delta-9-tetrahydrocannabinol (THC). Our results demonstrate a spatio-temporal patterning, with ventriculomegaly in THC-exposed embryos followed by a deceleration of brain growth in neonates that is sustained until adulthood, especially in females. We observed consistently impacted regions in both the cortex and subcortex, aligned with sex-dependent changes to social behavior in neonates and increased anxiety-like behavior in adolescents. Our results suggest prenatal THC exposure has a sustained sex-dependent impact on neurodevelopment that may persist into early adulthood.

## Introduction

As cannabis is legalized in more jurisdictions worldwide, it is also increasingly viewed as safe and natural,^1,2^ leading to greater use among pregnant people^3–5^ to manage symptoms related to morning sickness^6,7^ or anxiety.^8^ Importantly, the concentration of delta-9-tetrahydrocannabinol (THC) in readily-available whole plant cannabis and related products is also increasing.^9,10^ THC is the main psychoactive component of cannabis,^11^ and its use has been associated with long-term alterations in neurocognition, especially among adolescents.^12^ There is an urgent need to investigate how cannabis exposure before birth affects neurodevelopment throughout the lifespan. This information can inform the decisions of pregnant people, clinicians, and policy makers.

The endocannabinoid system (ECS) plays important roles in key developmental processes of the central nervous system (CNS), including embryo implantation, neuronal differentiation, and axon guidance.^13^ The two major endocannabinoids, anandamide (AEA) and 2-arachidonoylglycerol (2-AG), are tightly regulated over the course of development with AEA levels dropping before implantation and 2-AG levels slowly rising over the course of gestation.^13^ THC is a partial agonist at cannabinoid (CB)1 receptors and (less strongly) at CB2 receptors,^14^ which are present in glial cells and in the immune system. Introducing THC early in development could disrupt these processes and subsequently impact brain development. Investigations of THC exposure on neural stem cells report precocious neuronal and glial proliferation^14,15^ and increased stem cell proliferation and metabolism.^16^ Immunohistochemistry of embryonic human brain slices following prenatal THC exposure (PTE) demonstrate axonal rewiring and decreased CB1 bouton density.^17^ Finally, human developmental studies of prenatal cannabis exposure (PCE) show associations with mood disorders^18,19^ and reduced academic and professional achievement^20,21^ later in life.

Postmortem assessments can only characterize the impact of PTE cross-sectionally and in specific regions of interest, experimental strategies that may obscure variation in whole-brain neurodevelopmental patterning. To overcome these challenges, we used whole-brain, longitudinal, structural magnetic resonance imaging (MRI) in mice to noninvasively examine the impact of *in utero* THC exposure on brain development from the gestational period until adulthood.^22^ This approach is translatable to humans and reflects current trends in normative modeling.^23,24^ To better characterize the effects of PTE in our experiments, MRI was paired with behavioral assessments and postmortem analyses.^25–27^ We further examined sex differences given the significant evidence of sex-specific neurodevelopmental trajectories^22^ and responses to THC exposure both *in utero* and across the lifespan.^28^

We investigated the impact of THC exposure *in utero* in pregnant mice injected subcutaneously with 5 mg/kg of THC from gestational day (GD) 3-10 in three experiments: 1) with embryos extracted at GD 17 and imaged with *ex vivo* MRI; 2) in pups imaged on alternate days from postnatal day (PND) 3-10 with manganese enhanced MRI (MEMRI);^22^ and 3) in pups imaged on PND 25, 35, 60 and 90. In all cohorts, whole-brain, voxelwise volumetric differences were investigated with deformation based morphometry (DBM) and assessed with linear mixed-effects models (LMERs). In the first two experiments, the profile of glial cells and neurons were assessed with electron microscopy (EM) to identify dark, apoptotic, and dividing/daughter cells. Social behavior was assessed in the neonates with separation-induced ultrasonic vocalizations (USVs) while anxiety-like behavior (open-field test [OFT]) and sensorimotor gating (prepulse inhibition [PPI]) were assessed in adolescence. Together, the methods employed provide a thorough characterization of the impact of PTE through development to adulthood.

## Results

### PTE affects embryo volume and growth rate across the lifespan

The potential impact of PTE on offspring growth is of interest given previous findings of intrauterine growth restriction (IUGR)^29^ and the potential long-term impact of IUGR.^30^ Accordingly, we examined group differences in body volume or growth rate in each cohort (<u>Methods</u>). Embryos were extracted on GD 17 to acquire *ex vivo* high-resolution anatomical MRI images. Body volume was estimated from whole-body images by calculating the volume of a mask of the whole embryo (mm^3^). In the longitudinal experiments, pups were weighed prior to scans and perfusions. For each experiment, volume or weight was modeled with LMERs, accounting for litter size in the cross-sectional experiment (embryos) and litter size and subject ID in the longitudinal experiments (neonates and adults) with random fixed effects. For each statistical test, full models and results can be found in <u>Supplementary</u> <u>Results (SR) 5.</u> The experimental timeline for injections and outcomes can be found in Fig. 1. In the embryo experiment, a Null group was included, where dams were restrained without injection to control for the repeated injections stressor. All experiments included the THC group and a saline (Sal) control. The sample size per cohort following quality control of images was as follows, embryos: Sal, *n = 24* (10 females), THC, *n = 19* (10 females), Null *n = 27* (14 females); neonates: Sal *n = 26* (9 females), THC *n = 23* (13 females); adults: Sal *n = 24* (12 females), THC *n = 24* (13 females) (Fig. 2a). Comparing the Null group vs the Sal revealed a modest impact of the chronic injections on embryo body volumes, with Sal embryos being slightly smaller than Null embryos (*p = 0.009, standardized b=0.96*). Exposure to PTE further reduced embryo body volume as seen in the significant difference between the THC and Sal (*p = 0.02*, *standardized b= −0.90*) (Fig. 2b.i; SR 5.1).

**Fig 1.**
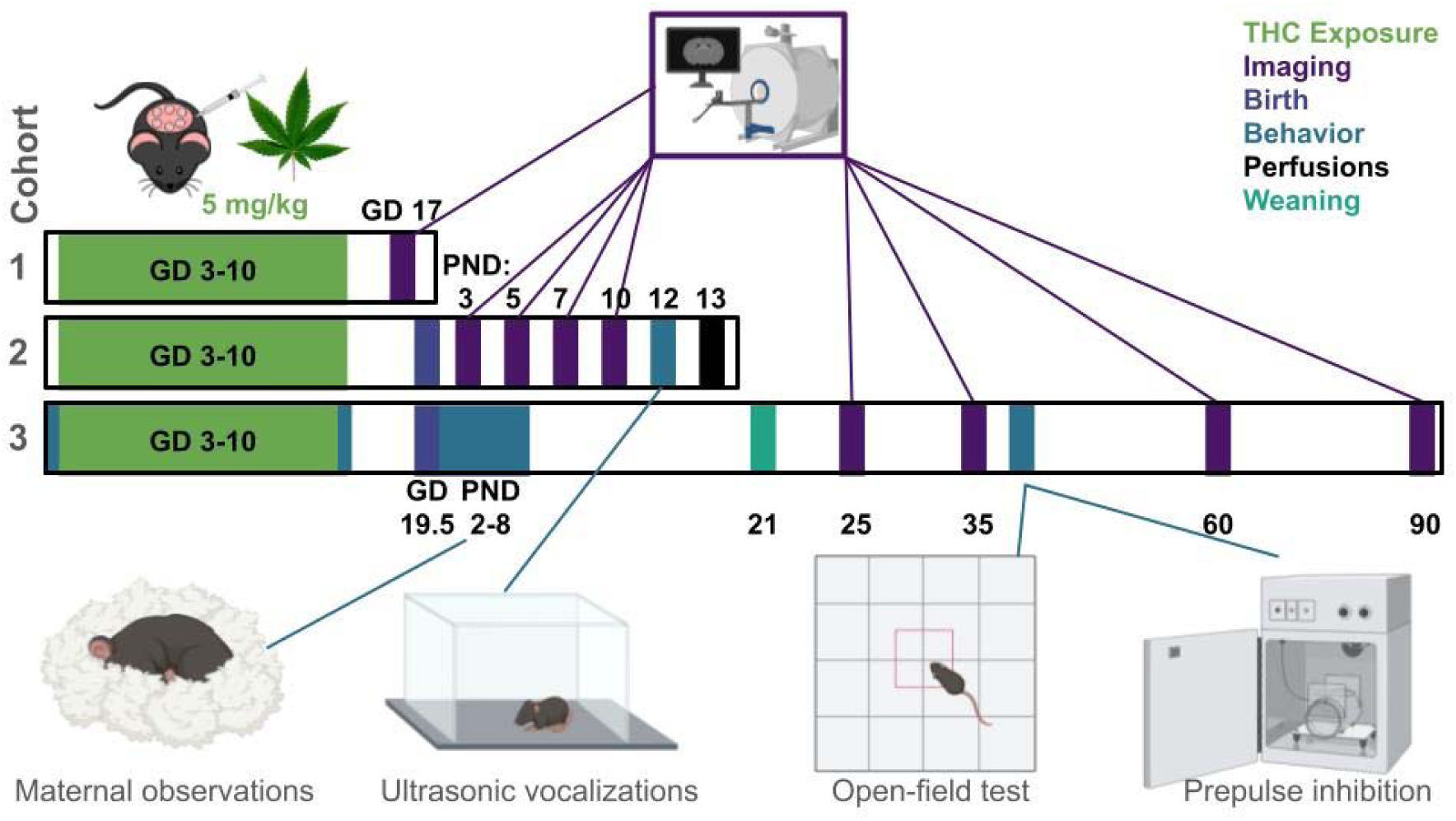
Experimental timelines. Timelines for the three experiments depicting s.c. injections of THC from GD 3-10. In cohort 1 (the embryo cohort), embryos were extracted and scanned at GD 17. In cohort 2 (neonate cohort), pups were born and scanned on PND 3, 5, 7, and 10. USVs were collected on PND 12 and pups were perfused on PND 13. In cohort 3 (adult cohort), nest quality was assessed before and after injections. Maternal behavior was assessed every other day from PND 2-8. Pups were weaned at PND 21 and scanned at PND 25, 35, 60, and 90.

**Fig 2.**
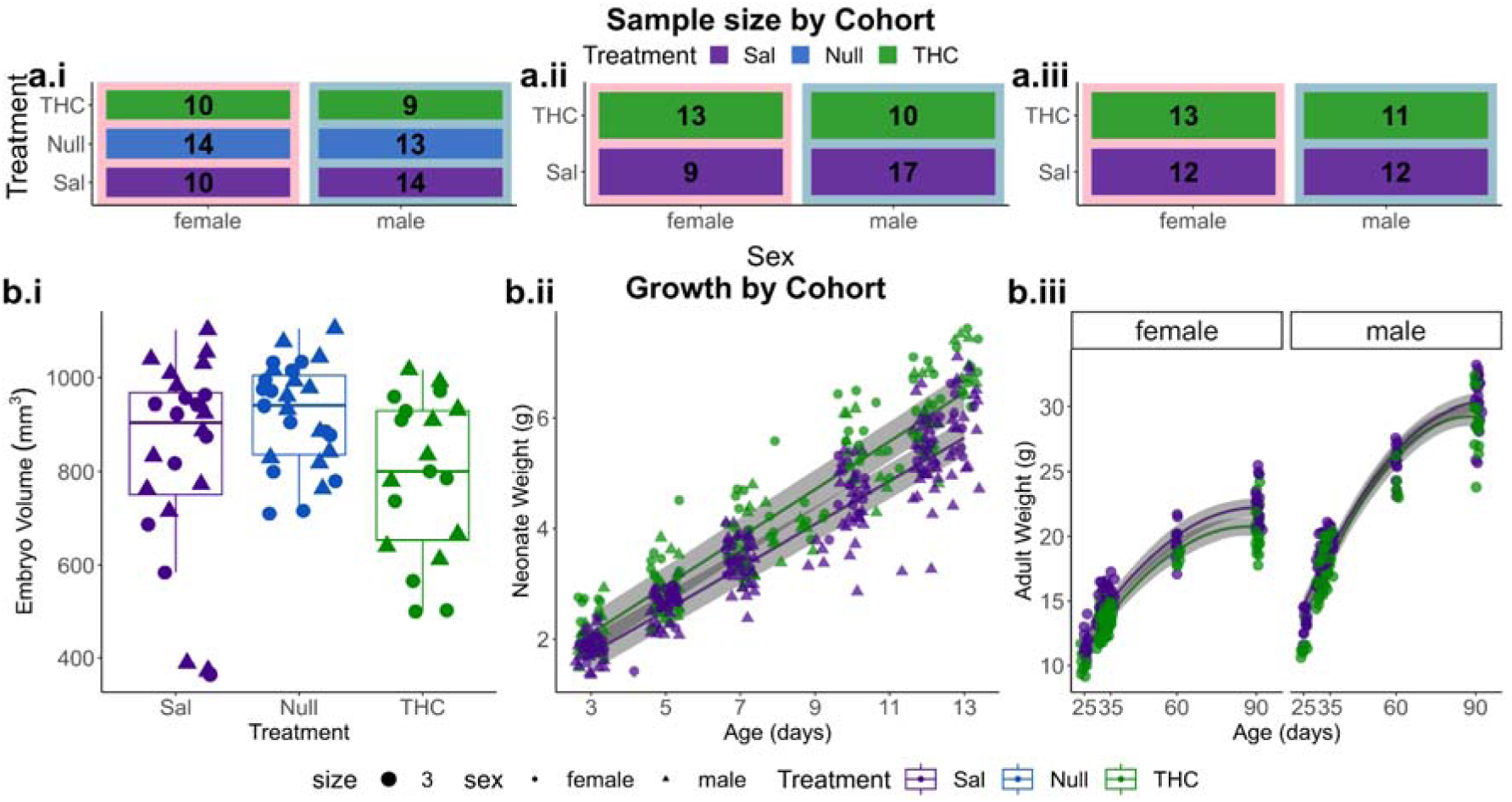
Sample size and offspring growth. a) sample sizes, following quality control across the three cohorts, embryos (left), neonates (middle) and adults (right). b) Embryo volume (left), neonate weight gain (middle) and adult weight gain (right). For embryos (b.i) Sal was the reference group to perform a single comparison of volumes across groups and maintain a consistent reference across experiments. Legend: size indicates the size of points. For sex, females are circles, males triangles. Sal is in purple, Null in blue, and THC in green.

Given the existing literature on sex differences and the ECS,^31^ we tested for an interaction between sex and treatment, but we observed no significant sex effect or sex-by-condition interaction. All pups gained weight over time (*p < 0.001*). However, while THC embryos were smaller *in utero*, THC neonates recovered over the first two weeks of life and gained weight more quickly than the Sal (age-by-condition interaction, *p = 0.003*) (Fig. 2b.ii, SR 5.2). No sex differences were observed in neonate weights (main effect or interaction with age or condition). We included non-scanned littermates (up to two per litter) in the neonate weight analysis, but there was no main effect of scanning on pups, indicating neither scanning nor anesthesia impacted growth. The emergence of the impact of PTE on sex-specific developmental trajectories became apparent at adolescence and adulthood (Fig. 2b.iii, <u>SR</u> <u>5.3</u>). We observed that males were heavier (main effect of sex; *p < 0.001),* and gained more weight over time (sex-by-age interaction; *p < 0.001).* PTE offspring also weighed less than controls even by the first timepoint (main effect of treatment; *p = 0.01)*. However, weight *change* was not impacted by PTE in the adolescent period (no treatment-by-age interaction). Nonetheless, as expected, all mice gained weight over time (main effect of age, modeled with both linear and quadratic terms; *p < 0.001*). There was no interaction between sex and treatment, or time and treatment, and the three-way interaction was not significant.

### PTE impacts brain volume and trajectories of growth across the lifespan

To assess brain volume, MRI was acquired in each cohort (scanning details in <u>Methods</u>). In embryos, the head was manually cropped to optimize image registration. Iterative unbiased DBM was used to create a consensus population average, employing image registration techniques across all input images and ensuring voxelwise correspondence for statistical analyses (<u>Methods</u>).^32^ Body volumes are likely related to brain volumes in embryos and should be accounted for as a nuisance term in models. In this study, we have already demonstrated a PTE effect on body volume; therefore the inclusion of body volume as a covariate introduces collinearity between the treatment and body volume. To account for this, we used a regression-with-residuals approach to correct for the mediating effect of body volume.^33^ This involved extracting the residuals from a model used to examine the effects of treatment on body volume and incorporating this value (representing body volume independent of condition) as a covariate in the brain volume model (<u>Methods</u>). Therefore, in the embryos, we used the condition-by-sex interaction and body volume residuals as fixed effects and litter size as the random effect in voxelwise LMERs. We then corrected with the false discovery rate (FDR). In cross-sectional analyses voxelwise volume was represented by relative Jacobian determinants capturing local differences, with global brain volume assessed separately (<u>Methods</u>, <u>SR 1.1</u>) and in longitudinal analyses voxelwise volume was represented by absolute Jacobians which account for both local and global volume differences, as with previous studies (<u>Methods</u>).^32,34^

THC embryos had larger bilateral lateral ventricle volumes and white matter regions, like the corpus callosum (main effect of condition significant, FDR 5%, heatmaps: Fig. 3a, peak voxels: Fig. 3b). Despite the modest effects of chronic injections on body volume, brain volume differences between the Null and Sal group were extremely small and focal in several anterior and medial regions (<u>SR 1.3</u>, 5% FDR). Neither sex nor a sex-by-treatment interaction were significant for brain volume.

**Fig 3.**
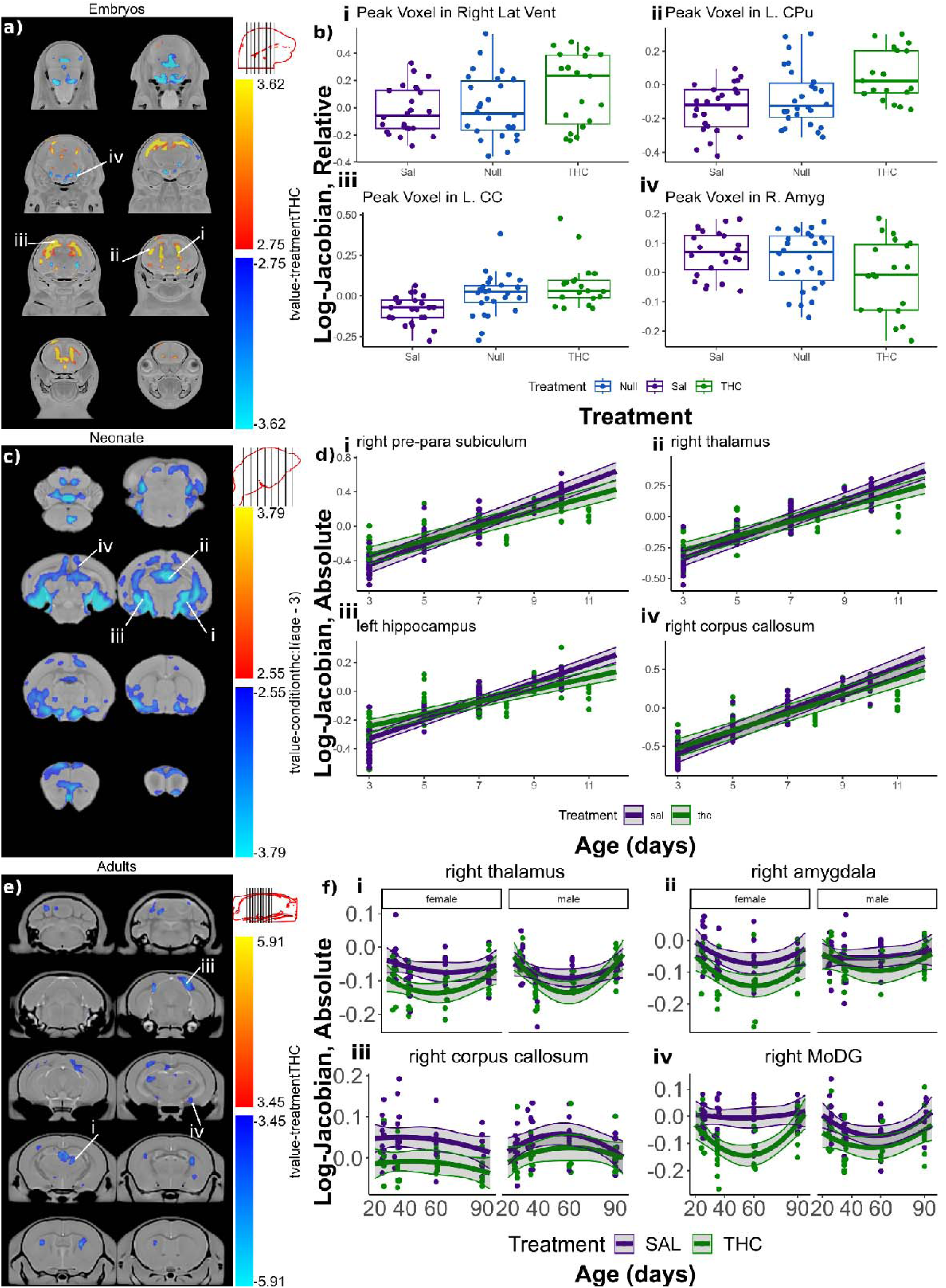
Brain results from MRI. a) Heatmap of volume differences between THC and Sal in the embryo cohort, thresholded FDR 5%. b) Peak voxels depicting volume differences in the embryo cohort. c) Heatmap of volume changes in THC pups compared to Sal controls (interaction with age), thresholded FDR 5% in the neonate cohort. d) Peak voxels depicting differences in rate of change in the neonate cohort. e) Heatmap of volume differences between THC and Sal controls (main effect), thresholded FDR 10% in the adult cohort. f) Peak voxels showing differences in THC group as well as between sexes (right amygdala [ii] not pictured).

In neonates and adults, we leveraged longitudinal *in vivo* structural MRI to examine trajectories of per-subject brain development, inspired by previous work in ascertaining brain development trajectories in both humans^35–37^ and rodent models.^22,26^ In the longitudinal cohorts (neonates and adults), we used a two-level DBM which is designed to capture individual growth curves while allowing comparison of these trajectories across groups (Methods).

We again used a regression-with-residuals approach to mediate the effect of weight (extracted from weight models). The LMERs for the voxelwise brain volume models (for neonates and adults separately) included fixed effects for condition-by-sex-by-age interactions and weight residuals, with subject ID and litter size as random effects. In neonates we found the linear age model best described the data, while in adults we observed that the addition of a quadratic effect of age (chosen with Akaike Information Criterion [AIC]) was a better fit for the voxelwise modeling (<u>Methods</u>).

In neonates we observed that PTE decreased growth rates in regions throughout the brain (condition-by-age interaction surviving 5% FDR, heatmap Fig. 3c). Significant variation in brain development includes brain regions such as the bilateral hippocampus, hypothalamus, amygdala, corpus callosum, cerebellum, striatum, ventricles, and thalamus (Fig. 3d). Unlike the neonatal cohort, the adults displayed no effects of THC-exposure on the rate of growth. Rather, there was a slightly less significant main-effect of condition, suggesting the overall effect was present at the first timepoint (10% FDR, heatmap Fig. 3e). The brain volume reductions that had emerged in neonates were still present in many regions in adulthood, including the hippocampus, hypothalamus, amygdala, cerebellum, inferior colliculus, corpus callosum, striatum, and thalamus (Fig. 3f), suggesting early volume reductions persist. Here, a sex-difference was observed (sex-by-condition interaction; FDR 10%). Overall, PTE contributed to smaller volumes, but THC males exhibited less volume reduction. Such regions include the bilateral olfactory bulbs, left cerebellum, medulla, piriform cortex, and arbor vita, as well as the right flocculus, striatum, and thalamus. Regions that showed the overall effect of THC have but the sex and treatment interaction included the left superior and right inferior colliculi, the left pons, and the right cingulate cortex.

### PTE impacts neonate calls and adolescent anxiety-like behavior

Behavioral changes have been characterized after PTE in both humans and animal models, and provide an important method to establish clinical relevance of experimental findings. Specifically, PTE and PCE have been highlighted as risk factors for anxiety-like behavior and social deficits.^19,38–41^ Human studies have also flagged a potential increase in psychosis-proneness in children prenatally exposed to cannabinoids, however this is a nuanced psychopathology that cannot be fully assessed in rodent models.^19,42^ Sensorimotor gating impairments have been demonstrated in humans with psychosis, although they are not specific to this symptom.^43^ Impairments to sensorimotor gating can be studied in rodent models with PPI, and mixed evidence of altered PPI has emerged following PTE.^39,44–47^

To determine if PTE altered relevant anxiety-like and social behaviors, mice in neonatal and adult cohorts underwent separate behavioral testing (<u>Methods</u>). Two days after the final scan, separation-induced USV were acquired in the neonate cohort to measure calls from the pup to their dam. During the USV test, each pup was separated from the dam for five minutes and the number and duration of calls were acquired. Variation in the call duration and frequency are reported as indicators for anxious and social behaviors.^48^ Distributions of calls were assessed with the hierarchical shift function^49^ (referred to as the shift function from hereon in), which uses permutation testing to determine how and by how much two groups of distributions differ (<u>Methods</u>).^32^ To assess the potential for sex-by-condition differences, subjects were divided by both condition and sex, with distributions of calls from the four sex-by-condition groups presented in Fig. 4a. The shift function revealed evidence for a sex-dependent effect of PTE. THC females exhibited deficits in social behavior, as they made fewer medium and long calls relative to Sal females. In contrast, THC males made more calls (specifically short and medium length calls), indicating an anxiety-like phenotype relative to Sal males. Finally, when examining the difference between Sal females and males as an overall sex effect, showing evidence for baseline sex differences, with females making more calls overall (Fig. 4b).

**Fig 4.**
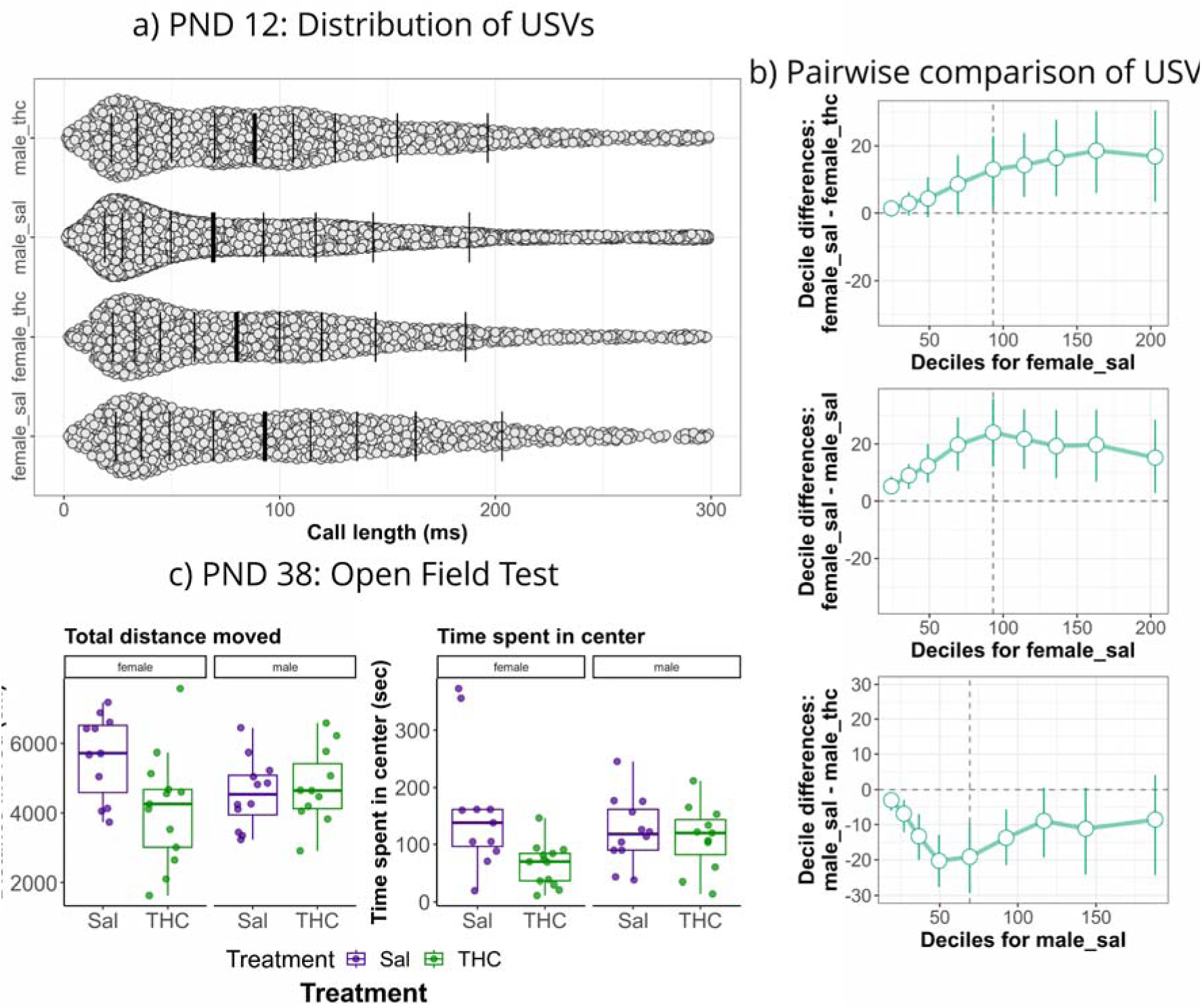
Behavioral results in embryos and adolescents. a) Distribution of calls pups made for dams with call length on the x-axis. From top to bottom, male THC group, male Sal group, female THC group, female Sal group. Each dot represents a call made from a pup to the dam. Vertical bars represent deciles and bold bars represent the median. b) Pairwise comparisons of USVs. Top: the difference between female Sal and female THC groups showing THC females make fewer long calls than female Sal. Middle: the difference between female Sal and male Sal indicating the main effect of sex, showing females make more calls over all. Bottom: difference between male Sal and male THC indicating THC males made more calls than male Sal. c) Left: differences in total distance moved for both condition and time spent in the center of the open field.

Behaviors were assessed in adolescence (PND 38-41, adult cohort), using the OFT to assay anxiety-like behaviors. We investigated group differences with total distance moved (overall locomotion), time spent in the center zone of the OFT (less time in center showing anxiety-like behavior^50^), and frequency of passes through the center zone (with fewer passes indicating anxiety-like behavior) using LMERs (fixed effects: condition-by-sex interaction, random effects: dam ID). We observed anxiety-like phenotypes in THC mice after Bonferroni corrections for multiple comparisons (Fig. 4c), with THC pups moving less compared to Sal controls (*p = 0.012*), spending less time in the center (*p = 0.003*), and making fewer passes through the center of the open field (*p = 0.005*). While none of the sex-effects survived correction for multiple comparisons (α = 0.016), there was trending evidence of anxiety-like behavior in females exhibiting hypolocomotion (*p = 0.047*) and male THC mice moving more than females (sex-by-treatment interaction; *p = 0.027*) and making more passes through the center of the open field (*p = 0.022*). Full results from the statistical models in <u>SR 5.4</u>.

PPI was used in the adolescent period to investigate sensorimotor gating. Percent PPI was calculated across trials, averaged at each prepulse intensity and assessed with LMERs (fixed effects: interaction between sex, treatment, and PPI level; random effects: subject ID and dam ID). Comparing THC pups to Sal controls, there was a trending effect consistent with impaired sensorimotor gating (decreased %PPI, p < 0.07, SFig 12, <u>SR. 3.2</u>), but no significant effect of or interaction with sex or 3-way interaction.

### PTE subtly impacts embryonic hippocampal cell division

Having identified the hippocampus (HC) as a region of interest from previous literature investigating the role of the ECS in network development,^51–53^ preliminary MRI findings from our neonates, and inspired by a previous study from our group linking MRI with postmortem assays,^32^ we used scanning electron microscopy (SEM) to image and quantify several categories of cells in the HC of both embryos and neonates (<u>Methods</u>). For the embryos and neonates, ∼4 samples per sex and condition were imaged at 25 nm resolution (full sample size in <u>Methods</u>). We first assessed the count of each cell type with LMERs (fixed effects: treatment and sex; random effects: subject ID; interaction between sex and condition were not considered due to sample size).

In embryos, we assessed dividing/daughter cells, dying cells, dark neurons, and dark glia, as microglia could not be differentiated from other glial cells at this stage. We only observed that more dividing/daughter cells were present in THC vs Sal embryos (Fig. 5a; p < 0.01), potentially consistent with previous observations of increased cell division following PTE,^15^ but surprising with the lack of volume change in this region at this timepoint. Because the distribution of dividing/daughter cells appeared different (Fig. 5b), we further tested the distributions with the shift function (Fig. 5c), finding significant differences between the Sal and THC groups, but not the Sal and Null groups. Compared to the Sal distribution the THC distribution was much wider with greater skewness, reflected in the long tail of samples showing high counts of dividing/daughter cells. This reflects a number of samples more greatly affected by the PTE.

**Fig 5.**
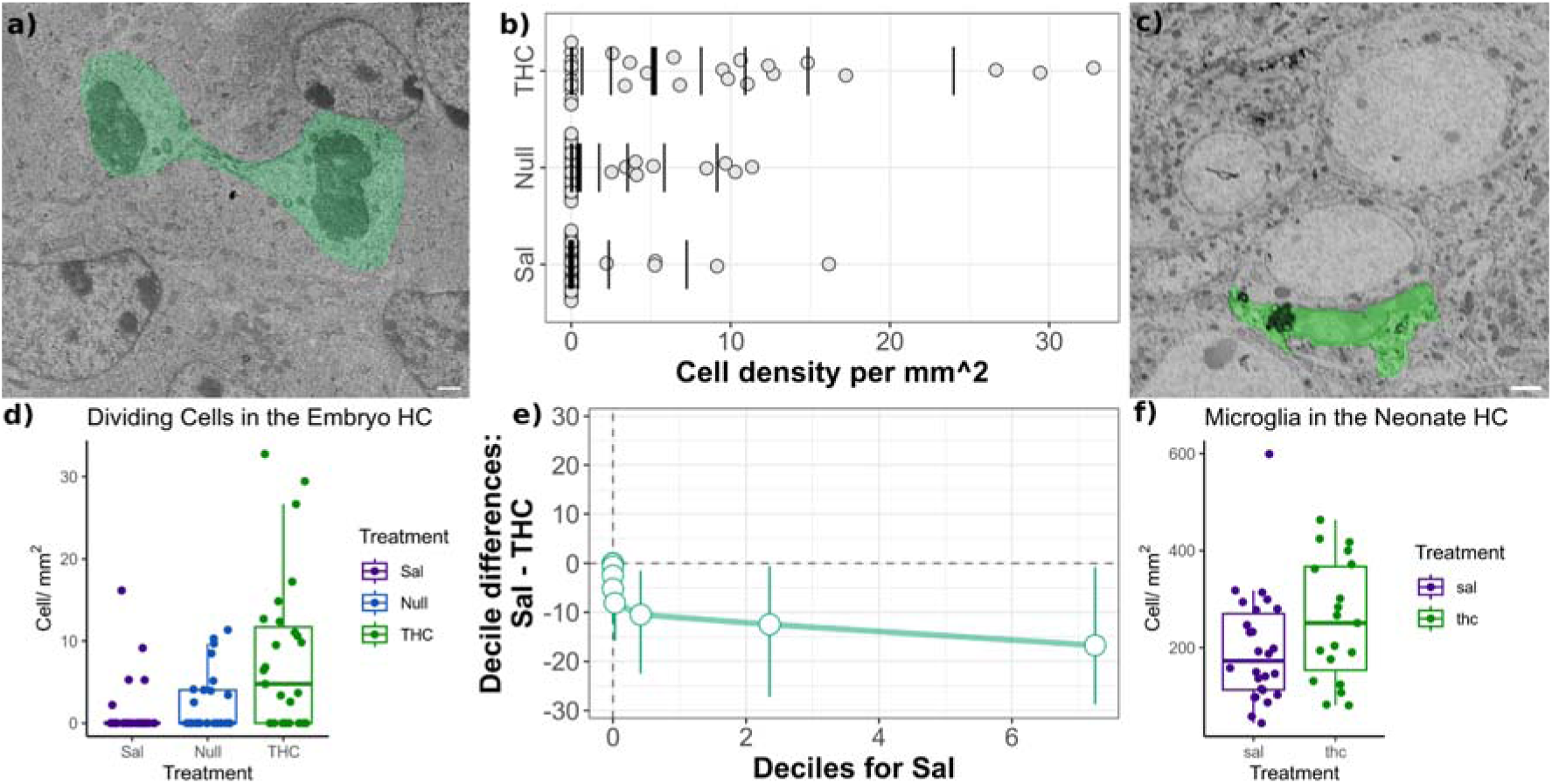
Electron microscopy in embryo and neonate hippocampi. a) Representative image of dividing cells in embryos (pseudocolored green, white scale bar = 1000 nm). b) Distribution differences between dividing/daughter cells in the embryo HC. c) Representative image of microglia in the neonate (pseudocolored lime, white scale bar = 2000 nm). d) Differences in dividing cell counts between Sal, Null, and THC embryos. e) Pairwise comparisons of differences between Sal and THC deciles in dividing/daughter cells in the HC. f) No difference between any examined cell types (microglia pictured) in the neonate HC.

In neonates, we examined apoptotic cells, dark neurons, dark glia, and overall microglia, as there was no evidence of dividing/daughter cells. There were no differences between the groups, suggesting changes in cell profiles may occur in embryonic periods but postnatal decreases in growth rate of the hippocampus are unlikely to be due to apoptosis or cell loss (Fig. 5d).

### Pregnancy outcomes and maternal behavior

To consider the effects of PTE on pregnancy-related outcomes and behaviors, we examined the impact of treatment on dam weight and weight gain, litter size, and two markers of maternal behavior: nest quality during pregnancy and time spent on nests in the early postnatal life. In brief, we pooled dam weight and litter-size data across experiments and found Null dams gained more weight than Sal controls (*p = 0.008*), and THC dams gained less weight than Sal controls (*p = 0.0002*), suggesting the THC had an impact above the effect of repeated injections (<u>SR. 4.1</u>). Regarding litter size, there was a main effect of treatment such that THC dams had smaller litters than Sal dams (*p = 0.016*; <u>SR. 4.2</u>). There was no impact of condition on nest building (<u>SR. 4.3</u>) or maternal behavior (<u>SR. 4.4</u>).

## Discussion

There is increasing urgency to understand the lifetime consequences of PTE. Despite clinical and preclinical studies that highlight the potential impact on behavior and specific brain processes, relatively little is known about the impact of PTE on trajectories of whole-brain development from the fetus to adulthood. Here we presented trajectories of growth and brain development with longitudinal data from late-gestation embryos to adulthood in mice.

Examining the whole-body volume of embryos and the weight and weight gain of offspring, we found that embryos exposed to THC had smaller whole-body volumes compared to Sal controls. The neonates showed catch-up growth, however from adolescence to adulthood there was a decrease in weight and weight gain which was more significant in females. These findings align with previous studies that demonstrate IUGR and low birth weight in both nonhuman animals and humans,^2,29,54–59^ as well as recent evidence of reduced weight-gain in PTE offspring.^60^ The catch up growth in the early postnatal life has rarely been documented in PTE models,^61^ however it is documented in other models of IUGR.^62^ This demonstrated reduction in fetal volume and alterations in weight gain throughout life could suggest changes to metabolism, which align with previously-published findings.^60,63^ For example, the ECS, and specifically PTE, have been demonstrated to play a role in altered liver and pancreas development, including dysmetabolism, and potentially impairments to glucose homeostasis.^64^

In the brain results, we see a distinctive pattern of ventriculomegaly in the embryos, followed by a reduced growth rate in neonates and sustained smaller volumes in the adults, especially the females. Although we were unable to use a consistent atlas across the embryo, neonate, and adult cohorts, we believe we have identified several regions as being consistently affected between GD 17 and PND 90 according to age-specific atlases. These included the caudate putamen, thalamus, olfactory bulbs, corpus callosum, amygdala, medulla, and hypothalamus. The ventricles were altered in embryos and neonates and the hippocampus was altered in neonates and adults. One possible mechanism of induced change is through CB1 receptors, which are rich in the basal ganglia, cerebellum, and hippocampus.^65^ Regions such as the hypothalamus and thalamus are thought to have relatively low levels of CB1 receptors, yet CB1 in these regions still play an important role in function.^66–68^ Other targets of interest include CB2 receptors, activation of which has been implicated in ventricle size in germinal matrix hemorrhage,^69^ G-Protein Coupled Receptor (GPR)55, which may modulate anxiety and play a role in neurodevelopment,^70^ Transient Receptor Potential (TRP) channels and Peroxisome Proliferator-Activated Receptors (PPAR), which can both be modulated by cannabinoids.^71,72^ The ventricles must also be affected by mechanisms beyond direct activation of receptors. The ventricles play a key role in the birth of new neurons, and it is possible that the ventriculomegaly is also associated with increased volume in periventricular regions in the embryos. CB1, CB2, and GPR55 have all been shown to promote the proliferation of neural precursor cells,^73^ however further investigations would be required to confirm these mechanisms. Finally, the large volume of the corpus callosum of THC-exposed embryos, followed by a decreased growth rate in neonates is consistent with evidence suggesting the role of early life THC exposure disrupting CB1 binding which contributes to errant spreading of developing corticofugal tracts.^17,74^ Additionally, developing white matter expresses high levels of CB1 receptors, potentially contributing to the early white matter changes observed.^75^

The behavioral results suggest alterations to social and anxiety-like behavior in neonates and adolescents respectively. Examining duration of USVs with the hierarchical shift function reveals basal sex-differences, as well as sex-dependent effects of condition. Typically, more or longer calls are interpreted as marks of distress or anxiety (as evidenced in the females compared to males and THC males compared to Sal males),^76^ but reduced calls have also been interpreted as impairments in social behavior (as evidenced in the THC females compared to Sal females).^77^ In general, evidence of separation-induced USVs following PTE are mixed.^78,79^ Without overinterpreting these results, it is evident there are sex-dependent alterations in social calling behavior in neonates. Anxiety-like phenotypes have been demonstrated in the OFT^80^ and the elevated plus maze^38^ following PTE, as well as in humans.^18^ These results align with our evidence from OFT, where THC-exposed female animals show decreased locomotion overall, as well as less exploration in the center of the open field. It is also possible that decreased locomotion was associated with worse motor performance, rather than anxiety-like behavior, as prenatal THC exposure has been implicated in poor motor coordination in rats.^81^ The PPI task did not indicate alterations in sensorimotor gating, with these null results consistent across several studies.^39,47^ One study did find that a single dose of THC in adolescence following PTE was sufficient to induce PPI alterations in males. This could suggest a priming effect of the PTE on sensorimotor gating.^82^

In general, we found limited effects in the EM outcomes. In neither the embryo nor the neonate experiments were there differences in the number of dark cells or dying cells in the hippocampus. We did see increased dividing/daughter cells in the embryo, consistent with studies that suggest CB1 receptor activation increases cell proliferation.^83^ The lack of other effects suggests that volume changes are not necessarily related to alterations in dark cell counts or apoptosis. However, potential functions of these cells, particularly microglia or dark cells were not assessed, which could influence changes in brain volume and behavior. The hippocampus was selected as it is known to be dense in CB1 receptors and has previously been investigated following alterations to cannabinoids during development.^84^ However, many of the hippocampal volume differences in MRI emerge only in neonatal periods. We believe our findings from our EM and early life provide some information to motivate further investigation at both finer grain anatomical and mechanistic levels in future studies.

Sex differences are an important and emerging focus of study in PTE, with robust evidence for sex effects following PTE in both human and animal model data.^31^ In our studies, we observed sex differences emerging in the adolescent/adult brain results, with both sexes exhibiting reduced volume but with some regions showing an altered trajectory in males compared to females. Additionally, in the behavioral results, there is evidence of sex-dependent effects in neonate USVs and trending sex effects (significant before correction for multiple comparisons) in the OFT during adolescence. There are several possible mechanisms that could contribute to sex differences following PTE, including differences in the ECS itself, differences induced by the effects of gonadal hormones, and sex-differences in the pharmacokinetics of THC metabolism.^31^ It is noteworthy that sex-differences in our MRI results only emerge after adolescence. It’s possible that PTE may have impacted organizational hormones prenatally which were then activated during puberty.^85^ Future studies should investigate the mechanisms underlying potential sex differences in PTE.

There are notable limitations to the approaches employed in these studies. First, we injected THC while most people inhale cannabis.^86^ Vape chambers have been increasingly used in the rodent PCE/PTE literature and have greater construct validity compared to injections.^87^ While evidence regarding the route of administration in pregnant people is lacking, there are reports that pregnant people may instead prefer to consume cannabis orally due to concerns about the potential effects of smoking on the fetus.^86^ There are differences in the pharmacokinetics of THC and its metabolites following injection or inhalation,^87^ as well as the amount of THC circulating in the maternal blood that crosses the placenta. One study compared subcutaneous injection of THC prenatally to exposure to vaporized THC, finding greater levels of THC in the fetal brain following injection than vaporization.^88^ Furthermore, when consumed orally, the bioavailability of THC is lower than when smoked, and peak absorption occurs more slowly.^89^ Overall, it is likely that the injected THC paradigm used here resulted in a higher concentration of THC in the fetal brain than would oral or vaporized cannabis preparations, which must be taken into account in interpreting the results. Likewise, in this study we injected THC alone, rather than a preparation including the other phytocannabinoids, such as cannabidiol (CBD).^90^ However, preparations of cannabis that are high in THC and low in CBD are increasingly popular,^91^ therefore it is still important to understand the effects of THC in isolation. Finally, the behaviors we tested were limited, as were the assessments performed on the data. For example, in the OFT, we did not measure thigmotaxis or rearing^92^ and in the USVs our software did not allow assessment of spectrogram typology of calls,^93^ which could provide additional information. Sample sizes several orders of magnitude larger would be required to reliably perform multivariate analyses such as partial least squares or canonical correlation analyses that relate brain and behavior.^94^

Overall, these studies provide one of the first developmental characterizations of the impact of PTE on development employing longitudinal MRI in rodents. Future studies could expand by investigating the underlying mechanisms for brain volume changes and sex differences or employing similar methods, such as longitudinal MRI in shorter periods in humans. In time, this work can contribute to a deeper understanding of how PTE impacts brain changes dynamically across the lifespan.

## Methods

### Animals and timed mating

All animals were housed at the Douglas Mental Health University Institute Animal Facility. Experiments were conducted in compliance with Facility Animal Care Committee regulations. Female and male C57BL/6J mice of breeding age (8 to 12 weeks old) were subject to timed mating breeding procedures, as described in <u>Supplementary Methods (SM) 1</u>. All females were nulliparous and males were mated for up to two litters. The day of an observed plug was considered GD 0. Animals were kept on a 12:12 light dark cycle (8am-8pm) and had *ad libitum* access to food and water.

### Drug preparation and injection

Dams were randomly assigned to Null (*n = 8*), Sal (*n = 20*), or THC (*n = 18*) condition, depending on the experiment (Null included only in embryo cohort). Null dams were restrained in a scruff hold and the injection area was touched with a gloved finger-tip, but did not receive injections. Because embryo brain results indicated only subtle differences between Null and Sal offspring (<u>SR 1.3</u>), the Null group was excluded from the neonate and adult experiments. In the Sal and THC groups respectively, dams received subcutaneous injections of vehicle (1:18 cremophor:saline) [Sigma Aldrich], or THC solution (1:1:18 THC:cremophor:saline) [Cayman Chemicals].^95^ Manipulations were performed daily from GD 3-10 between 9:00-11:00 am.

A research license and import permit for THC were obtained from Health Canada, and 2 g of THC were purchased from Cayman Chemical Company in a formulation of 10 mg/ml in ethanol. Ethanol was dehydrated, and THC was prepared in a solution (<u>SM 2</u>) as described previously.^25^

The dose of THC injected was 5 mg/kg. This dose has been characterized in both prenatal THC models,^39–41,88,96^ as well as adolescent exposure models.^25,38^ While it has been suggested that this dose represents moderate human gestational exposure,^39^ it can be challenging to accurately assess and model human cannabis use during pregnancy. The dose used here roughly corresponds to a dose of 27.6 mg/kg in humans, which is likely a moderate-high dose, as discussed in <u>SM 2.</u> The injection volume for both THC and Sal dams was 0.01 ml/g. Dams were weighed daily at the time of each injection and their weight recorded for analysis.

### Offspring outcomes and timeline

#### Embryos

Embryo extraction was conducted following previously published methods.^32^ On GD 17, pregnant dams were euthanized by live cervical dislocation. The uterus was immediately dissected and placed into a petri dish of chilled Phosphate Buffered Saline (PBS). In turn, each embryo was removed to a dish of warm PBS to stimulate blood flow, uterine tissue and the yolk sac were carefully removed, and a small piece of the yolk sac was collected for genotyping the SRY gene to determine embryo sex (Transnetyx, Memphis, Tennessee, USA). The umbilical cord was cut to encourage blood drainage and mice were placed in a vial with warm PBS. The vial was placed alternately between a shaker to help exsanguination and an incubator to maintain PBS temperature until blood had cleared. After approximately 10 minutes, each embryo was checked and either returned to warm PBS for further drainage or transferred to a vial containing 4% paraformaldehyde (PFA) as a fixative and 2 mM Gadoteridol (Gado) (ProHance; MRI contrast agent; Bracco Imaging S.p.A) for 1 week at 4°C. After 1 week, embryos were transferred to a long-term storage solution of 2% Gado and 0.02% sodium azide 1x PBS solution at 4°C until scanning. Gado was used as a contrast agent for *ex vivo* MRI, as it is absorbed into tissue and reduces the T1 and T2 relaxation times,^97^ enhancing the contrast, especially with long scan times.^98^ Up to 2 male and 2 female pups from each litter were sent to the Mouse Imaging Centre, Hospital for Sick Children, Toronto (scan and image processing details in <u>SM 3.1</u>) to minimize potential litter affects.^27^

#### Neonates

On PND 1, litters were culled to 6 pups. On PND 2, 4, 6, and 9, 24 hours before each scan (details in <u>SM 3.2</u>), dams were injected with 0.4 mmol/kg dose of 30 mM manganese chloride (MnCl_2_) solution, a contrast agent that pups ingest while nursing.^22^ MnCl_2_ shortens the T1 and T2 relaxation times, increasing the contrast of images^99,100^ and had been used in previous work from our group.^26,101^ Of note, MnCl_2_ has been shown to impact development, transiently reducing weight gain of pups and brain volume (differences peaking about −10% before normalizing for both) discussed further in <u>SM 3.3</u>.^102^ Despite these side effects, the method has been used successfully in developmental studies before.^22^ On PND 3, 5, 7, and 10, 4 pups per litter, 2 males and 2 females where possible, were scanned. Before each scan, pups were examined for developmental milestones including physical (weight, incisor eruption, fur development, eye opening, pinnae detachment and auditory canal opening), sensorimotor (Vibrissa placing response, ear twitch, and auditory startle response), and motor (surface righting reflex and grasp reflex) metrics adapted from previous publications.^103–106^ A full description of the tasks and timeline is presented in <u>SM 4.1</u>, <u>SR 2.1</u>. On PND 12, pups were separated from their dams for 5 minutes and USVs they made during this time period were recorded to analyze the frequency and duration of calls. On PND 13, pups were weighed and sacrificed by transcardiac perfusion, and their brains were extracted for postmortem assays.

#### Adults

Maternal behaviors were assessed in the adult cohort only, as embryo extraction was a terminal procedure and assessments during the neonatal period may interfere with maternal behavior. The quality of nests dams built during the dark period was assessed at two timepoints, the night before the first injection (night of GD 2-3) and the night after the last injection (night of GD 10-11). On PND 1, all pups were weighed and litters were culled to six. Observations of maternal behavior were made on PND 2, 4, 6, and 8 at 9:00 (one hour after lights on) and 21:00 (one hour after lights off). On PND 21, pups were sexed and weaned. On the two days prior to the first scan (PND 25), pups were handled for a minute to habituate them to handling. The second scan was acquired on PND 35. OFT was acquired on PND 38, and PPI was acquired on PND 41. Finally, the third scan was acquired on PND 60, and the fourth on PND 90, scan details in <u>SM 3.4</u>. Pups were sacrificed on PND 91.

### Image acquisition and processing

#### Embryos

Embryo samples were shipped in Styrofoam boxes packed with ice and ice packs. In advance of imaging, each sample was removed from the vial with contrast solution, blotted dry, and placed in 13-mm diameter plastic tubes filled with a proton-free susceptibility-matching fluid (Fluorinert FC-77, 3M Corp. St. Paul, MN), described previously.^32^ Images of the whole embryo body were acquired on a multichannel 7 Tesla (T) MRI scanner with a 40-cm-diameter bore (Varian Inc.). A custom-built 16-coil solenoid array was used to acquire images of up to 16 samples simultaneously, facilitating long scan times (14 hours) overnight with an ultrahigh resolution voxel size of 40-µm isotropic.^32,107^ Further detail is provided in <u>SM 3.1</u>. After image reconstruction, they were preprocessed to enable statistical analysis, as in previous work from our group.^32,108^ First, a mask was created and corrected manually to exclude peripheral tissue and bubbles that were present in some images to enable extraction of the full embryo image. Each image was quality-controlled by two independent raters (L.C. and A.P.) to assess whether the sample was misoriented in comparison with a reference image and whether the sample had been distorted or damaged in transportation (e.g., compressed and therefore no longer morphologically or anatomically accurate). Fourth, an in-house preprocessing pipeline was used to correct for intensity inhomogeneities and denoise the images. Further information about each step is included in <u>SM 3.1</u>.

Following pre-processing, images were registered through an unbiased DBM pipeline built in-house with a Python wrapper around the antsMultivariateTemplateConstruction2.sh from the ANTs environment.^109^ In brief, an unbiased model building process is performed on all images to create a study-specific average and ensure voxelwise correspondence across all images. Specifically in the case of the embryos, the images were further manually registered to the average produced by the DBM before a second complete pass of the DBM. Precise details can be found in <u>SM 3.1</u>.

Within the DBM pipeline, in the first stage, affine transformations are used to create a group average of all input images accounting for bulk changes in size and shape. In the second stage, nonlinear transformations are used to iteratively estimate the residual differences in contrast profiles among images and transform images to minimize these differences. Together, these iterated affine and nonlinear transformations produce a minimal deformation field which maps each individual subject image to the average at the voxelwise level. The Jacobian determinant of the warp field can then be calculated to provide voxelwise estimates of volumetric difference relative to the average representation, with a value from from 0 to 1 indicating a smaller volume and a value >1 indicating a larger volume relative to the average.^110–112^ Statistical comparisons can be performed on a voxelwise logarithmic transformation of the derived Jacobians as this better conforms to assumptions required for the use of parametric statistical analyses. The relative Jacobians explicitly comprise only the volumetric differences modeled by the nonlinear part of the deformations and explicitly remove the contribution of the global linear differences (attributable to differences in total brain/body size).^32,110^ The relative Jacobians were smoothed with a Gaussian kernel at 80-µm full-width-at-half-maximum to better conform to Gaussian assumptions for downstream statistical testing.^32^ A mask of the peritoneal cavity and organs (including heart, lungs, and liver) was made on the average and transformed to each of the input images for an exploratory analysis examining volume differences among organs (<u>SR. 1.2</u>). Then, the head was cropped in all images and head-only images were used for a third and final DBM, thereby optimizing registration for the brain. Further detail on the DBM and QC can be found in <u>SM 3.1</u>. Brain regions impacted by PTE were identified by comparison to an atlas.^113^

#### Neonates

The day of the scan, pups were removed from the cage one at a time. They were weighed, assessed for age-appropriate milestones (e.g. incisor eruption starting PND 5 and fur development on PND 7), and anesthetized with isoflurane (induced at 5%, maintained at 2% in 100% oxygen 1 L/min during the scan). Pups were fit into foam beds carved for each age and placed into the scan bed. A nose-cone was fitted over the animal’s snout to deliver isoflurane, and the mouse was heated with warm air delivered to the bed, as to adult mice.^25,34^ Following the first scan timepoint on PND 3, pups’ paws were micro-tattooed with India ink for identification. After the scan, pups were rubbed with bedding to assure their dam would accept them back into the nest, returned to their home-cage, and placed on a heating mat.^22,102^

After images were reconstructed, they were converted to the NIFTI file type, quality controlled, and preprocessed, with full details reported in <u>SM 3.2</u>. Images were processed with the same in-house longitudinal two-level DBM pipeline as used in later-life studies^25^ and cross-sectionally in embryos. In contrast to the embryos, the pipeline was used over two-levels of registration to account for the longitudinal design, wherein an unbiased average was first constructed for each subject over time and then across subjects. The final step of the DBM pipeline involved post-processing the consensus deformation fields to produce Jacobian determinants which encode the voxelwise volumetric difference from each input scan to the consensus population average (ANTs toolkit). Registrations were quality controlled and, because of the great changes in brain size and morphometry between PND 3 to 10, we observed that there were many failed registrations, especially in the cerebellum. We improved these registrations, which are critical to the performance of the pipeline, by applying a manually developed brain mask on the initial unbiased average using the Display software available in minc-toolkit-v2. This mask was transformed to the individual input images with ANTs tools (antsRegistration and antsApplyTransforms) using transforms produced in the first run of the DBM and then manually corrected.^109^ Subject-level warp masks were quality controlled and manually corrected where necessary and then used to extract the brain from the background of the image (ImageMath). The DBM pipeline was run again and results quality controlled to ensure all images were well-registered.

#### Adults

Before each scan, the mouse was weighed and anesthesia was induced with 3.5% isoflurane in 100% oxygen at 1 L/min. Pups were transferred to the scanner, where anesthesia was maintained at 1.5% isoflurane, administered in an 80% air, 20% oxygen mixture at 1 L/min. Offspring were restrained, and warmed by hot air to maintain body temperature. On PND 25, 35, 60, and 90, structural MRI was collected for 30 minutes. After the first scan, mice were identified with an ear notch, and after each scan they were allowed to recover for 5 minutes on a heating pad under the home cage.

All mice were scanned on the same scanner as the neonates. The structural scan was a fast low-angle shot sequence with the following parameters: 3D gradient echo sequence, TR = 20 ms, TE = 4.5 ms, flip angle = 10°, field-of-view = 18.0 × 16.0 × 9.0 mm, matrix size = 180 × 160 × 90, number of averages = 2, total acquisition time = 10 minutes 48 seconds, isotropic resolution = 100 µm^3^.

As in previous studies^25^ and the neonates, after images were reconstructed, they were converted to NIFTI files, quality controlled, and preprocessed, with further details reported in <u>SM 3.4</u>. Following preprocessing, we employed the same longitudinal, two-level DBM pipeline as for the neonates to obtain individual Jacobian determinants within the adult population average space and proceeded to voxelwise comparisons (<u>SM 3.4</u>). Compared to the neonate mice, adult mice did not require any manual corrections as the changes in brain volume were more subtle between PND 25 and PND 90 than between PND 3 and 10.

### Behavioral analyses

#### Neonates

##### Ultrasonic vocalizations

USVs were assessed on PND 12, 48 hours after the final scan. Due to technical malfunctions (distributed across conditions and sex), data were available from a restricted sample of the scanned animals. Therefore calls were assessed two ways. First, results were analyzed in the pups exposed to MnCl_2_ <u>SR 2.4</u>, with the following sample size: Sal female - 7, THC female - 15, Sal male - 15, ThC male −12. Then, this sample was combined with additional pups that were unexposed to MnCl_2_ but otherwise exposed to an identical experimental paradigm with the following sample size: Sal female - 5, THC female - 2, Sal male 6, THC male - 8. Impact of MnCl_2_ on call duration was assessed with an LMER with pup ID and litter as random effects and no significant difference based on MnCl_2_ exposure. USVs were assessed with procedures that have been previously reported.^114–116^ The dam was removed to a clean cage and pups were tested one-by-one. The home cage was placed on a heating pad and in turn each pup was isolated in the behavioral testing room, placed in the apparatus of the Noldus UltraVoxTM system (Noldus Information Technology, Leesburg, VA). The plexiglas box was also placed on a heating pad. After USVs were collected, the mouse was weighed and any milestones collected before it was placed back in the home cage. Pups were rubbed with bedding material following the final pup’s testing and the dam was replaced in the cage.

#### Adults

##### Open-field test

On the day the OFT was conducted, mice were brought to the testing room 30 minutes after lights on and allowed to habituate for 30 minutes. Up to four opaque, gray OFT boxes (45 cm^3^) were placed in the field of view of an overhead camera. A mouse was gently placed in each open field box and allowed to explore for 15 minutes while video was recorded. Mice of the same sex were tested together. To assess behavior, videos were analyzed with the software Ethovision XT12 (Noldus Information Tech Inc., Leesburg, VA). The bottom of the open field captured on video was conceptually divided into zones including a center zone (40% of the area) and the outer edge and corners. The mouse was tracked automatically, and the total distance traveled was analyzed, as well as the duration of time in and the passes through the center zone. All tests were completed within the first 4 hours after lights on.

##### Prepulse inhibition

As with OFT, on the day PPI was conducted (2 days after OFT), mice were habituated to the testing room for thirty minutes. PPI to acoustic startle was measured with commercially produced startle chambers (San Diego Instruments, San Diego, CA). Details of the chambers have been previously reported^26^ and are available in the <u>SM 4.2</u>. Mice were gently placed in the holder in each PPI chamber and the pre-programmed experiment was run.

### Scanning electron microscopy

#### Embryos

After imaging with MRI, embryos were shipped to the University of Victoria, British Columbia, for SEM imaging. Upon arrival, the brains were extracted from the sample and postfixed with 3.5% acrolein in phosphate buffer (100 mM) (pH 7.4) overnight at 4°C. Following the quality control of MRI-images, samples with high quality images were selected for EM-analysis (Null, n = 8 [2 female, 6 male], Sal, n = 8 [4 female, 4 male], THC, n = 7 [4 female, 3 male]. Brains were sectioned to 50 µm sagittal slices with a VT1200S vibratome (Leica Biosystem), and stored at −20 LJ in cryoprotectant (30% glycerol and 30% ethylene glycol in PBS [50 mM] [pH 7.4]). Slices that contained the hippocampal formation (coronal section 18-19, sagittal 3-5, horizontal 6),^113^ were processed with osmium tetroxide osmium (OTO) fixation, dehydrated, and polymerized in Durcupan^TM^ ACM resin (Sigma-Aldrich) as previously described.^117^ The hippocampus was excised from the resin and glued onto a separate block of resin before being sectioned into approximately 70 nm ultrathin sections with a Leica Artos 3D ultramicrotome. After slicing, the ultrathin “ribbons” of 3-5 sections were captured on silicon wafers (EMS) at 8-10 µm intervals. Tissue was loaded into a Zeiss Crossbeam 350 SEM and three-four images per embryo were acquired at 25 nm of resolution in x-and-y with the SE2 detector at a voltage of 1.4 kV and current of 1.2 nA.^32^

Images from the three treatment groups (Null, Sal, and THC) and both sexes were analyzed blind to the condition by H.A.V. with the QuPath software (v0.5)).^118^ Each image was then manually examined and the number of dying cells, dark glia, dividing/daughter cells and dark neurons were calculated. Cell type was determined by ultrastructural properties. We previously described these properties for dark neurons and dark glia—briefly, dark cells were determined by their dark cytoplasm and fainter nuclear heterochromatin pattern, with dark neurons being generally larger than glia and containing an apical dendrite or, occasionally, shrunken and deformed but possessing a pyknotic nuclei, whereas dark glia were smaller, occasionally had smaller, irregular protrusions and had clumping in their faint heterochromatin pattern.^32^ Dying cells were defined by a pyknotic or fragmented nucleus, an incomplete nuclear envelope with nuclear condensation (including “necklace” patterning)^119^ or an accumulation of autophagic vacuoles, as described previously. Dividing/daughter cells were identified by nuclear chromatin patterning, and were mostly found to be either in prophase, prior to nuclear envelope breakdown, or more commonly in the final stage of mitosis/telophase with cell constricted between it in cytokinesis.^32,120,121^ Area around the ROI was manually traced and the area was calculated to determine density of cells of interest and total cells.

#### Neonates

One day after the behavioral tests, on PND 13, pups were given an intraperitoneal injection of ketamine, xylazine, and acepromazine at a dose of 0.1 mL/100 g before being transcardially perfused. Perfusion was performed using 1x Phosphate Buffered Saline (PBS) solution with heparin, followed by 4% PFA. Brains were extracted from the skull and post-fixed in the same paraformaldehyde (PFA) solution for 24 hours. After 24 hours, the brains were transferred from the PFA-PBS solution to a solution of PBS and 30% sucrose for cryoprotection.

Following perfusion, *ex-cranio* neonate brains were shipped to the University of Victoria, British Columbia, for EM imaging, like the embryos. The postfixation, processing, and imaging procedure was identical to that of the embryos, except the neonates were sectioned coronally. The sample size included: Sal, n = 7 [3 female, 4 male], THC n = 5 [3 female, 2 male].

In the neonates microglia and dark microglia were counted versus dark glia overall, as they are more able to be distinguished at this time point without subsequent immunostaining. Microglia and dark microglia were identified with ultrastructural properties as previously described. Briefly, microglia were identified by their typical heterochromatin pattern^122^ and dark microglia were identified by their electron dense appearance, loss of the microglial heterochromatin motif and ultrastructural markers of cellular stress.^123–125^

### Statistical analyses

#### Pregnancy outcomes following PTE

All statistical tests were performed in the R programming language, version 3.5.1, with packages lme4_1.1-33, lmerTest_3.1-3, rogme_0.2.1, and RMINC_1.5.3.0. Data were pooled across experiments to assess the impact of PTE on pregnancy outcomes in terms of dam weight gain, and litter size. We examined the impact of treatment on weight gain in the dams with an LMER (lme4::lmer and lmerTest::summary), including a fixed effect of the interaction between treatment and GD, litter-size as a covariate, and dam ID as a random effect. The impact of treatment on litter size was also assessed with linear models.

#### Embryos

##### Magnetic resonance imaging

First we examined the impact of the interaction between condition and sex on total body volume. As it is zero-bounded, volume was scaled to have zero mean and unit standard deviation to comply with assumptions of LMERs. Including multiple pups from a single litter violates assumptions of statistical independence among individuals, and litter is considered a nesting variable.^26,27,126^ Therefore, LMERs were employed to include a random effect of litter, as well as coil position, since multiple samples were scanned on the same coil.^126^ Instead of using coil number (1-16), coils were categorized by their position as either “corner” (4 coils), “edge” (8 coils), or center (4 coils). The fixed effects of the model included an interaction between condition and sex.

To examine the impact of condition and sex on local volume differences, we employed identical LMERs on the log-transformed relative Jacobians from the DBM, resulting in a single model per voxel. Because IUGR has been shown to impact brain volume,^127^ we sought to control for the effects in our model. As previous literature and evidence from this study suggest THC exposure impacts body volume,^29,58^ simply including volume in the model as a covariate could introduce co-linearity between condition and volume. Therefore, we employed a regression-with-residuals method to assess the mediating effect of body volume on the relationship between condition and brain volume.^33^ The residuals from the total body volume model were extracted and included as a covariate in the brain volume model. Thus, our fixed effects included the interaction between treatment and sex, and the body-volume residuals, and the random effects were litter size and coil category. To correct for multiple comparisons we used the FDR, where a threshold from 5% indicates up to 5% of significant voxels are false positives. As supplementary analyses to the main results, we examined the impact of treatment on total brain volume (TBV), <u>SR 1.1</u>. Using the brain mask we calculated volume under the mask for each individual in R (RMINC::anatGetAll). We compared the conditions with an LMER, examining the impact of sex and treatment and including litter as a random effect.

##### Electron microscopy

For each of the four EM metrics assessed (dying cells, dividing/daughter cells, dark glia, and dark neurons), an LMER was run examining the main effect of treatment and sex and including animal ID as a random effect. For significantly different comparisons, the hierarchical shift function was used to examine distributions in the data.

Descriptive measures, such as means or medians, may not accurately represent differences between groups, but the shift function (rogme_0.2.1::shifthd pbci) can be leveraged to compare distributions of data.^49^ In this approach, the distributions of data are mapped between groups, with, for example, each distribution comprising the number of cells per group. The median and deciles of the distributions are calculated per condition (Sal and THC). Deciles are estimated with the Harrell-Davis quantile estimator (HD).^49,128^ The differences between the distributions are quantified; specifically, with the shift function it is estimated how and how much one distribution must be shifted to match another one.^49^ Then, 95% confidence intervals per decile comparison are calculated with bootstrap resampling. To achieve this, the original data are resampled with replacement within condition 2,000 times. The number of resampled observations is determined by the number of iterations (in this case 2,000) multiplied by the number of observations per condition. For each iteration, the HD is calculated for the resampled distributions and the difference between deciles between the two resampled groups are calculated. After all resamples are complete, a p-value representing the significance of the difference between groups for the observed deciles is calculated by examining the proportion of resampled cases where the differences between groups is more extreme than the observed differences between groups. This value ranges from 0 to 1, with 0 indicating no resampled cases are more extreme than the observed difference (very significant observed difference) and 1 indicating that the resampled cases are consistent with the observed difference (not at all significant observed difference). The value is multiplied by 2 to generate a two-sided p-value. Finally, Hochberg’s correction for multiple comparisons is applied.^129^ Deciles where the confidence interval does not cross 0 indicate deciles that significantly differ between groups.^49^

#### Neonate

##### Weights

LMERs were used to assess the impact of treatment and scanning on pup’s weight, scaled to have 0 mean and unit standard deviation. The weight model included an interaction between condition, age, and sex with scanning status (whether the pup was scanned or not) as a covariate, and littersize and animal ID as random effects.

##### Magnetic Resonance Imaging

As in the embryo experiment mass univariate LMERs were run on the Jacobian determinants, however in this experiment, the absolute instead of relative Jacobians were primarily assessed. Absolute Jacobians, as opposed to relative Jacobians, include affine components of registration, encoding changes in total brain volume which can be useful for identifying gross volumetric changes over time. In embryos this was represented in the separate total brain volume analysis (<u>SR 1.1</u>). Relative Jacobian differences for neonates can be found in <u>SR 2.2</u>. Again, because of the impact of treatment on weight and weight gain, the regression-with-residuals approach was used to control for the impact of treatment on weight gain without introducing collinearity into the model. The residuals from the pup weight model were extracted and included as a covariate in the brain volume model. Thus, the fixed effects included the interaction between treatment, sex, and age, as well as the weight residuals, and the random effects were litter size and animal ID. To correct for multiple comparisons we used FDRs. Voxels were automatically labeled using RMINC tools (mincFindPeaks, mincLabelPeaks) from a PND 7 atlas adapted from an adult atlas and used in previous neonatal experiments,^114,130^ as PND 7 represents the mid-point between the included ages.

##### Ultrasonic vocalizations

First, to exclude background noise, calls were filtered for those between 5 and 300 ms.^48,114^ Then, to analyze differences in frequency of USVs of different lengths, we compared the distribution of calls, as previously with the shift function.^114^ To examine the interaction between sex and treatment, data were separated into four groups: sal female, sal male, THC female, THC male. Comparisons in distributions were calculated between each of the groups. Supplementary examinations of the calls made by scanned vs unscanned pups were also included following the same statistical approach (<u>SR 2.3</u>).

##### Electron microscopy

As in embryos, for each of the EM metrics assessed (dying cells, dark glia, and dark neurons, and total microglia), an LMER was run examining the main effect of treatment and sex and including animal ID as a random effect. Because there was no evidence for dividing/daughter cells, this comparison was excluded.

#### Adult

##### Weights

Weights of the developing mice were acquired on scan days (PND 25, 30-35, 60, and 90) and the day they were sacrificed (PND 91). Trajectories of weight were assessed with LMERs, with fixed effects including the interaction between treatment, sex, and age (modeled as a quadratic function), and random effects of subject ID and litter.

##### Magnetic resonance imaging

A mass univariate, whole-brain, voxelwise analysis was conducted with tools from the RMINC package (version 1.5.3.0) on log-transformed absolute Jacobian determinants produced by the DBM. Like with the neonates, relative Jacobians are included as supplementary analyses (<u>SR 3.1</u>) As in previous studies, our evidence suggests THC exposure impacts offspring weight gain, and weight could impact brain volume, therefore we employed a regression-with-residuals method to control for how body weight might mediate the relationship between condition and brain volume. The residuals from the pup weight model were extracted from that model and included as a covariate in the brain volume model. Therefore, fixed effects included: the interaction between treatment, sex, and the quadratic effect of age, as well as the weight residual, and random effects included: subject ID and dam ID. Multiple comparison corrections were conducted with the FDR.

##### Open field test

Total distance moved, duration of time in the center zone, and frequency of passes through the center zone were compared with LMERs. Fixed effects included the interaction between treatment and sex, with litter as a random effect.

##### Prepulse inhibition

Percent PPI was calculated and averaged across trials for each prepulse intensity with the following equation:

(startle response − prepulse response)/startle response ∗ 100

Percent PPI was assessed with LMERs, with fixed effects including the interaction between sex, treatment, and PPI level, and random effects included subject ID and mom ID.

## Supporting information

Supplemental methods and results

