## Supplemental methods and results for "Impact of prenatal delta-9-tetrahydrocannabinol exposure on mouse brain development: a fetal-to-adulthood magnetic resonance imaging study"

Supplement

### Supplementary Methods

#### 1. Animals

All C57BL/6J mice were housed in the animal facility in conventional ventilated cages with a 12-hour light cycle (8:00-20:00) with *ad libitum* access to food (standard rodent chow) and water. Female and male mice of breeding age (8-12 weeks) underwent timed mating. One male and one female were placed into a new cage in the late afternoon (15:00-17:00). The following morning (8:00-9:00), the male mouse was removed and the female was checked for the presence of a seminal plug and weighed on a scale with a readout to the tenths of grams. Mice were paired for one night only to improve the accuracy of the timing of mating. When a seminal plug was observed, this was considered GD 0. If no plug was observed, a minimum of nine days elapsed before the female mouse was weighed again to check for pregnancy attributed weight gain (>2g), and if she was still of a breeding age, timed mating was attempted again. If the female was pregnant, she was sacrificed or her offspring used for other experiments/the next generation of parents. Males were used multiple times.

#### 2. Drug Preparation

To eliminate the potential influence of trace amounts of ethanol from the formulation, the solution of THC (Cayman Chemical Company, Annarbor, Michigan, USA) in ethanol was dehydrated in a Universal vacuum system UVS 400/supervac SCIIOA to obtain a highly viscous, dehydrated THC. Because THC is lipophilic, it must be dissolved in a 1:18 vehicle of cremophor (Sigma-Aldrich) and saline. A stock solution of 0.5 mg/mL of THC in solution was prepared to obtain an injection volume of 0.01 mL/g at a dosage of 5 mg/kg. The stock solution was then aliquoted to 1.5 ml Falcon tubes and frozen to prevent THC degradation in light and heat. The vehicle solution of 1:18 cremophor:saline was also prepared, aliquoted, and frozen, to avoid any confounding effects of cremophor or freezing.

Preparations of dosages for 2.5, 5, and 10 mg/kg of THC in stock solution were validated in a separate cohort of adult mice (both sexes) with gas chromatography mass spectrometry to ensure that the expected dosage was being injected, and the confirmatory results have been previously published ^1^. The injected dosage of THC was 5 mg/kg per day in dams, administered subcutaneously from GD 3-10. The Human Equivalent Dosage (HED) can be converted from the murine dosage with Equation S1.

Eq. S1: HED(mg/kg) = AnimalDose(mg/kg) ∗ AnimalKm/HumanKm

where Km is a species-specific constant, which is a factor that normalizes by the body surface area ^2^. In converting from mice to humans, this equation becomes:

Eq. S2: HED(mg/kg) = 5 ∗ 3/37

which solves to 0.405 mg/kg for a human.^2^

For a 68 kg person, this would result in a dosage of 27.54 mg of THC, which is difficult to interpret in the context of smoked cannabis or use during gestation. While the exact answer is not available, there are certain published statistics about cannabis preparations that can aid in the assessment of dosage. First, the average THC concentration of joints (cannabis cigarettes) seized in Northern California in 2019 was 14%, which corresponds to approximately 140 mg of THC.^3,4^ Second, donated joints from adult cannabis users in Barcelona contained a median of 6.56 mg of THC, but a range from approximately 3-75 mg of THC.^5^ Third, among daily users, participants in the United Kingdom self-reported their cannabis use, estimating that they smoked about 1 gram each day, with a THC concentration of about 9%, using about 0.42 g of dried cannabis in each joint, which would result in joints containing roughly 37.8 mg of THC.^6^ These studies suggest great variability among cannabis users and much higher consumption of THC in preparations from daily users than from occasional recreational users. Together they provide evidence that our dosage is reasonable for a moderate-heavy dosage of THC for a daily user, although, to date, no study specifically assesses THC consumption in pregnant populations.

#### 3. Structural MRI

##### 3.1 *Embryos*

The structural scan was a T2-weighted image acquired with a 3D fast spin echo sequence using a cylindrical k-space acquisition, with TR/TE=350/12 ms, echo train length=6, two averages, field-of-view 20 mm x 20 mm x 25 mm, matrix size=504 x 504 x 630/^7–9^ The 4-by-4 array for 16 sample acquisition introduces geometric distortions as the samples are placed at varying distances from the isocenter of the gradients in the magnet. Specifically, the farthest samples from the isocenter are most susceptible to artifacts.^8^ Therefore, corrections are applied during image reconstruction to assure high-quality images are obtained regardless of position in the array discussed in great detail in:^8^ 1) the amplitude of each echo is intensity corrected with a Wiener deconvolution, 2) each readout is aligned and phase-matched, 3) phase correction was predicted and removed from k-space prior to reconstruction, 4) where even and odd k-spaces were combined, they were aligned with translation and averaged prior to reconstruction.^8^

In preprocessing the reconstructed image, the first step was mask creation. The Advanced Normalization Tools (ANTs) and minc-toolkit-v2 ecosystems were used to first clamp negative intensities from the image (mincmath command with -clobber and -clamp flags) then to create an Otsu mask (ThresholdImage command). Full commands can be found in publicly available code. Once each mask was created, the Display command from the MINC Toolkit library was used to visualize each image with the mask as a label superimposed on it (<https://bic-mni.github.io/>). The masks were then corrected manually by either L.C. or A.P. to exclude any tissue outside of the bounds of the embryo body. Corrected masks were used to estimate embryo volume with the in-house script, collect_volumes.sh, a wrapper around Insight Toolkit’s itk_label_stats, available open-source online (<https://github.com/CoBrALab/minc-toolkit-extras/blob/master/collect_volumes.sh>). Then the masks were multiplied by the original image to result in an image with a clean boundary at the edge of the embryo body. Next, each image was quality controlled in comparison to a reference image, a GD 18 average from a previous experiment.^9^ The command register from the MINC Toolkit was used to compare orientation, and images were assigned a value of 0 if they were misoriented or 1 if they were properly oriented. They were also assessed for whether they were deformed (1) or not deformed(0). If an image was noted as misregistered, “tags” were created indicating similar parts of the image and the reference (such as the mouse snout, back of head, and top of head). Then linear transformations were applied to the individual image to align it with the reference. Images that were noted as deformed were removed from future analyses. In total, 14 images were removed for being deformed, (Null, N = 1 [female = 0], Sal, N = 4 [female = 4], THC, N = 9 [female = 4]). Finally, an in-house pipeline used ANTs tools to preprocess the individual scans, including denoising with adaptive nonlocal means (minc_anlm) and N4 bias-field correction.^10^

A DBM pipeline was used to align the images and create a study-level average. First, all images were aligned with affine transformations that preserved parallel lines in the images (translations, rotations, scales, and shears). Finally, iterative nonlinear transformations attained precise anatomical alignment between images in an automated fashion, similar to previous studies in adult animal brains,^1^ as well as embryo studies in our laboratory.^9^ After the initial DBM, images representing the transforms from the individual scans to the study average were visually inspected, and if the registration failed, the original input image was visually inspected to identify the cause of the misregistration. One major cause of mis-registration was that some of the embryos are more curled, or c-shaped, from the tip of the nose to the tip of the tail. In order to assist the registration to the rest of the population, the c-shaped embryos were identified. Again, tags were created with the MINC tool “register” between the input image and the study-average produced from the DBM. The tags were converted to an LSQ12 transform file (tagtoxfm -lsq12) and the transformation was applied to the input image (mincresample). The LSQ12-transformed images were then re-registered to the study-average and tags between the two images were produced again. These tags were converted to a thin-plate spline non-linear transformation (tagtoxfm -tps), and the transformation was applied to the LSQ12 images. Another DBM was conducted, replacing the c-shaped images with the TPS-transformed images. This model-build resulted in 5 registration failures, which were excluded from the final analysis (Null, N = 2 [female = 0], Sal, N = 3 [female = 2], THC, N = 0). Finally, to optimize registration for the brain, the head was cropped from each image and another run of the DBM pipeline was performed, as per above. A cubic mask was made with the SimpleITK package in the Python programming language identifying the head of the average from the last DBM. This was then transformed to each of the input images using the transforms produced by the DBM (ANTS:antsApplyTransforms) and the bounding box was QC’d. Then, images were converted to mincs (nii2mnc) and the mask was applied as a bounding box to crop the images (minc-toolkit::autocrop –bbox). These cropped head images were used as inputs for a final DBM.

##### 3.2 Neonates

First, images were converted to NIFTIs with the BrkRaw module tool, *brkraw tonii.*^11^ Next, they were quality controlled by two independent raters (L.C. and A.P.). Each scan was examined for motion and artifacts and given a score from 1 to 5, with 1 being the best and 5 being unusable. Scans that received a 1 or 2 from both raters were included, those that received a 4 or 5 were excluded, and scans that received a 3 or where there was disagreement were examined again for a consensus to be reached. Next, all scans were pre-processed with an in-house pipeline using ANTs and minc-toolkit-v2 tools to denoise each image with adaptive nonlocal means (minc_anlm) and apply an N4 bias-field correction.^10^ Following preprocessing, in-house software was used to conduct a twolevel model-build, first registering all images within a subject, and then across subjects to create a population average (https://github.com/CoBrALab/optimized_antsMultivariateTemplateConstruction). In each level of the model build, all images were aligned with affine transformations that preserved parallel lines in the images (translations, rotations, scales, and shears). Then, iterative nonlinear transformations attained precise anatomical alignment between images in an automated fashion as before.

##### 3.3 MnCl_2_ in neonates

In the course of the study, the potential site-specific effects of postnatal MnCl_2_ exposure on Sal and THC pups was estimated. To this end, four additional litters (two Sal, two THC) were acquired unexposed to MnCl_2_. Twenty-four hours prior to each scan, the dam was injected with Sal, but otherwise the experimental design was identical to the full experiment. Unexposed pups were included in USV experiments, and information on their weight and brain growth examined elsewhere.^12^

##### 3.4 Adults

The scanning protocol for adults was adapted from previous neurodevelopmental studies in another center.^13^ All mice were scanned on a 7T Bruker BioSpec with a 30 cm bore magnet with AVANCE III electronics, equipped with a cryogenically cooled surface coil. The T1-weighted scan protocol had the following parameters: 3D gradient echo sequence, TR = 26 ms, TE = 5.37 ms, flip angle = 37°, field-of-view = 17.1 × 14.8 × 11.1 mm, matrix size = 190 × 164 × 123, number of averages = 2, total acquisition time = 23 minutes 19 seconds, isotropic resolution = 90 µm^3^.

Again, images were converted to NIFTIs with the BrkRaw module tool. Next, they were quality controlled by two independent raters (L.C. and K.B.). Each scan was examined for motion and artifacts and given a score from 1 to 5, with 1 being the best and 5 being unusable. Scans that received a 1 or 2 from both raters were included, those that received a 4 or 5 were excluded, and scans that received a 3 or where there was disagreement were examined again for a consensus to be reached. Next, all scans were pre-processed with an in-house pipeline using ANTs and minc-toolkit-v2 tools to denoise each image with adaptive nonlocal means (minc_anlm) and apply an N4 bias-field correction, as before. Following preprocessing, the same DBM pipeline reported in the neonates was used to generate the population average and Jacobian Determinants, without the need to extract the brains, as in the neonates.

#### 4. Behavior

##### 4.1 *Pup milestones*

Pup milestones were adapted from previous research.^14–17^ They were designed to be as minimally invasive as possible and, therefore, assessed only on days when the pups were separated from their dams for scanning or behavioral assessment. First, pup weight was assessed at PND 3, 5, 7, 10, 12, and 13. The anogenital distance was assessed at PND 3, when pups were sexed. Incisor eruption was assessed at PND 10 and 12. Fur development and pinnae detachment was assessed on PND 3 and 5. Eye and auditory canal opening was assessed on PND 12 and 13. Vibrissa placing response was assessed by tickling the whiskers with a Q-tip and marked as present if the mouse turned its head towards the Q-tip. It was assessed on PND 5, 7, 10, 12, and 13. Ear twitch was assessed by tickling the pinnae of the ear on PND 10 and marked as present if the ear twitched. Auditory startle response was assessed on PND 7, 10, 12, and 13, by clapping the hands and assessing whether the mouse flinched in response. Surface righting was assessed on PND 7. Mice were held supine and, when released, a stopwatch was started. If the mouse turned prone in less than 1 second, surface righting was considered present. Grasp reflex was assessed on PND 5, 7, 10, 12, and 13. The middle section of a Q-tip was brushed against the forepaws of the pup, and if it curled its paws around the bar and exerted some force when the bar was drawn away, grasp was assessed as present. For all behaviors and physical milestones, the pup was assessed for first appearance of the milestone, with the exception of weight which was assessed on all specified days.

##### 4.2 *PPI*

Acoustic startle chambers from San Diego Instruments consist of a Plexiglas, cylindrical chamber (8 cm diameter, 16 cm long) set on a Plexiglas base in a sound-attenuating chamber. A speaker is located in the ceiling 24 cm above the animal which provides background noise (70 dB) as well as acoustic stimuli. Beneath the cylindrical chamber where the mouse is placed is a piezoelectric accelerometer which detects and transducts motion accompanying acoustic startle. A microcomputer using a commercial software package from SR-LAB was used to control pulse parameters, digitize, and record the stabilimeter readings. After animals were placed in the cylindrical chambers, they underwent 5 minutes of habituation to the background noise before undergoing a total of 50 trials (5-30 s intertrial duration). The first 8 and final 7 trials measured startle magnitude to a stimulus (50 ms 120 dB) without prepulse, and the middle 35 trials the startle tone was presented alone or preceded by a 30 ms prepulse ranging from 3-15 dB above the tone (3 dB increments: 73-85 dB). The prepulse varied randomly between trials with 5 trials presented per prepulse stimulus. The average startle response was derived from the 100 1 ms readings taken after onset of the startle stimulus.

### Supplementary Results

#### 1. Embryos

##### 1.1 *Total brain volume*

TBV between groups was assessed with an LMER, where the fixed effects included an interaction between treatment and sex and the random effect included the litter. The equation for the model is as follows:

TBV model: Y = 𝝱_0_ + 𝝱_1_SexM + 𝝱_2_conditionTHC+

𝝱_3_conditionNULL + 𝝱_4_conditionTHC:sexM + 𝝱_5_conditionNULL:sexM + **b**_1_litter + **ϵ**

Y= outcome measures (i.e. TBV in mm^3^); 𝝱_i_= fixed effect coefficient; 𝝱_0_ = equation intercept; **b** =

random predictor; **ϵ** = random error; **:** = interaction;

conditionTHC= prenatal THC group relative to SAL as reference; conditionNULL = prenatal NULL group relative to SAL as reference; SexM = male sex relative to female as reference

There was no effect of either sex nor treatment on TBV (SFig 4).

##### 1.2 *Organ volume*

While there was no impact of the Null condition compared to Sal, there was both a main effect of THC and the interaction between condition and sex for THC significant from 10% to 5% FDR. The main effect of sex showed a trending effect from 15% to 10%. The main effect of THC showed clusters of voxels in the liver (medial and left lobes, larger in THC), and the lungs (mixed results, SFig 5). The interaction showed reduced volume in the right liver of THC males compared to Sal females, not pictured. The equation for the organs was as follows:

Organ model: Y = 𝝱_0_ + 𝝱_1_SexM + 𝝱_2_conditionTHC+

𝝱_3_conditionNUL + 𝝱_4_conditionTHC:sexM + 𝝱_5_conditionNULL:sexM + **b**_1_litter + **ϵ**

Y= outcome measures (matrix of voxel-wise measurements); 𝝱_i_= fixed effect coefficient; 𝝱_0_ = equation intercept; **b** =

random predictor; **ϵ** = random error; **:** = interaction;

conditionTHC= prenatal THC group relative to SAL as reference; conditionNULL = prenatal NULL group relative to SAL as reference; SexM = male sex relative to female as reference

##### 1.3 *Sal vs Null, local brain volume*

There were several focal anterior, medial regions (near the preoptic area) that showed a smaller volume in the Null group compared to the Sal controls (5% FDR), seen in SFig 6. The equation for the brain volume was as follows:

Brain volume model: Y = 𝝱_0_ + 𝝱_1_SexM + 𝝱_2_conditionTHC+

𝝱_3_conditionNUL + 𝝱_4_conditionTHC:sexM + 𝝱_5_conditionNULL:sexM + **b**_1_litter + **ϵ**

Y= outcome measures (matrix of voxel-wise measurements); 𝝱_i_= fixed effect coefficient; 𝝱_0_ = equation intercept; **b** =

random predictor; **ϵ** = random error; **:** = interaction;

conditionTHC= prenatal THC group relative to SAL as reference; conditionNULL = prenatal NULL group relative to SAL as reference; SexM = male sex relative to female as reference

##### 1.4 *Embryo absolute Jacobians*

To complement the TBV analysis and the relative Jacobian analysis, we compared absolute Jacobians with the same model as the relative Jacobians in the main body of the paper. There was a main effect of treatment at 5% FDR, such that THC animals showed large volume in the ventricles and smaller volume in the hypothalamus, much as in the relative Jacobians (SFig 7).

#### 2. Neonates

##### 2.1 *Pup milestones*

Milestones were assessed with LMERs to examine whether condition, sex, or scanning impacted the age of achievement of a milestone (with dam_ID as a random effect). There were no significant differences, with the exception of the achievement of vibrissa placing response, which the females achieved slightly earlier than the males (*p = 0.04*; not pictured). The equation for first appearance of milestone was as follows:

Milestone model: Y = 𝝱_0_ + 𝝱_1_SexM + 𝝱_2_conditionTHC+

𝝱_3_scannedYes + 𝝱_4_conditionTHC:sexM + **b**_1_litter + **ϵ**

Y= outcome measures (e.g. age of achievement of Vibrissa placing); 𝝱_i_= fixed effect coefficient; 𝝱_0_ = equation intercept; **b** =

random predictor; **ϵ** = random error; **:** = interaction;

conditionTHC= prenatal THC group relative to SAL as reference; scannedYes = whether pups were scanned, with yes relative to no; SexM = male sex relative to female as reference

##### 2.2 *Neonate relative brain volume differences*

Relative Jacobians capture local nonlinear differences without the gross differences across the entire brain (linear transformations). For the relative Jacobians modelled with the same model as absolute Jacobians presented in the main manuscript, neonates showed a main effect of age, age*condition, and weight residuals thresholded from 5% to 1% FDR. The age*condition interaction results are presented in SFig 8. Regions in blue indicate a smaller growth rate in the THC pups over time in specific regions, over-and-above the reduced whole-brain growth rate. Regions in warm colors indicate focal areas of local volume increase over time in THC-exposed pups despite generally smaller growth rates in these regions.

##### 2.3 *Impact of scanning on USVs*

A final comparison between scanned and un-scanned pups, regardless of condition, with the hierarchical shift function reveals un-scanned pups make more medium-length calls than scanned pups, potentially indicating anxiety-like behavior among the scanned pups, SFig 9.

##### 2.4 *Impact of MnCl_2_ on USVs*

To control for the impact of MnCl_2_ on USVs, condition and sex of USV results were assessed only in the subsample exposed to MnCl_2_ (as this was the larger subsample) with the hierarchical shift function. This revealed the same direction of results as the full sample, but with less significance, as is expected from a smaller sample (SFig 10).

#### 3. Adults

##### 3.1 *Adult relative brain volume differences*

For adults, relative volume differences were assessed with the same model as absolute differences in the main body of the manuscript. Unlike absolute Jacobians, relative Jacobians did not show a main effect of THC, but they did show a condition-by-sex interaction with regions in the cerebellum showing larger volume in THC males (against the female Sal reference group; SFig 11).

##### 3.2 *Prepulse Inhibition*

As expected, with increasing prepulse level percent PPI increased as well (*p < 0.001*). There was a trending effect (*p < 0.07*) of THC exposure such that THC exposed mice exhibited decreased percent PPI, indicative of impaired sensorimotor gating, SFig 12. The equation for PPI was as follows:

PPI Model: Y = 𝝱_0_ + 𝝱_1_SexM + 𝝱_2_conditionTHC+

𝝱_3_PPILevel + 𝝱_4_conditionTHC:sexM + 𝝱_5_conditionTHC:PPILevel + 𝝱_6_conditionTHC:PPILevel:sexM + **b**_1_litter +**b**_1_subject + **ϵ**

Y= outcome measures (Percent PPI); 𝝱_i_= fixed effect coefficient; 𝝱_0_ = equation intercept; **b** =

random predictor; **ϵ** = random error; **:** = interaction;

conditionTHC= prenatal THC group relative to SAL as reference; SexM = male sex relative to female as reference; PPILevel = prepulse level (in decibels)

#### 4. Maternal and Pregnancy Outcome

##### 4.1 *Impact of treatment on dam weight and weight gain*

To establish PTE’s impact on pregnancy outcomes, we pooled data across the three cohorts to examine weight gain and litter size. Dam weight gain (LMER; w/ fixed effects: treatment and GD; random effects: dam ID). The Null dams gained more weight than the Sal controls (*p = 0.008;* treatment-by-age interaction), indicating a potential impact of repeated injections. Further, THC dams gained less weight than Sal controls (*p = 0.0002*), suggesting the THC had an impact above the effect of repeated injections (SFig 1). There was also a main effect of treatment, such that Null dams weighed less than Sal controls at GD0 (*p = 0.001*) despite our best efforts to randomize dams across groups. The equation for dam weight gain was as follows:

Dam weight gain model: Y = 𝝱_0_ + 𝝱_1_GD + 𝝱_2_conditionTHC+

𝝱_3_conditionTHC:GD + **b**_1_subject + **ϵ**

Y= outcome measures (dam weight); 𝝱_i_= fixed effect coefficient; 𝝱_0_ = equation intercept; **b** =

random predictor; **ϵ** = random error; **:** = interaction;

conditionTHC= prenatal THC group relative to SAL as reference; GD = gestational day

##### 4.2 *Impact of treatment on litter size*

The impact of treatment on litter size was assessed with a linear model (SFig 2). There was a main effect of treatment indicating that THC dams produced smaller litters than Sal dams (*p = 0.016)*. There was no difference between the Null and Sal groups. The equation was as follows:

Litter size model: Y = 𝝱_0_ + 𝝱_1_conditionTHC + 𝝱_2_conditionNULL +

**ϵ**

Y= outcome measures (litter size); 𝝱_i_= fixed effect coefficient; 𝝱_0_ = equation intercept; **ϵ** = random error; **:** = interaction;

conditionTHC= prenatal THC group relative to SAL as reference; conditionNULL= prenatal NULL group relative to SAL as reference

##### 4.3 *No impact of treatment or injections on nest quality*

In order to assess whether changes to maternal care and behavior were sufficient to explain developmental differences postnatally, we examined the quality of nests built pre-or-post-injections as well as maternal behavior postnatally in the adult cohort since in the neonatal experiment the litter was involved in scanning.

When examining nest quality with an LMER (fixed effects: session-by-treatment interaction; random effects = dam ID), there was no evidence for an effect of the injections as the session (pre-or-post-injection) did not significantly impact the quality of nests built. Additionally, there was no evidence that PTE impacted the quality of nests built, and no interaction between the two variables (SFig 2). The equation was as follows:

Nest quality model: Y = 𝝱_0_ + 𝝱_1_session + 𝝱_2_conditionTHC+

𝝱_3_conditionTHC:session + **b**_1_subject + **ϵ**

Y= outcome measures (nest quality); 𝝱_i_= fixed effect coefficient; 𝝱_0_ = equation intercept; **b** =

random predictor; **ϵ** = random error; **:** = interaction;

conditionTHC= prenatal THC group relative to SAL as reference; session = pre-or-post injections

##### 4.4 *No impact of treatment on postnatal maternal behavior*

Maternal behavior was assessed on PND 2, 4, 6, and 8, for 10 minutes, one hour after the light-cycle began and one hour after the dark-cycle began ([Methods](https://docs.google.com/document/d/17p5RXiOwHLbzGvhzgvDu6Ofucr71055I6io2GQCR7OA/edit#bookmark=id.4sb35srbryd5)). We used LMERs to assess both duration of time spent on the nest as well as frequency of passes over the nest (fixed effect: treatment-by-PND-by-time of day interaction;random effect: dam ID. Duration was calculated as percent duration, (duration of time spent on nest/total time *100). There was a trending effect of PND (*p = 0.07*), indicating that dams spent more time on the nest as the days went by, however there was no main effect of treatment or time of day, and no interaction among the variables (SFig 3a). In terms of frequency of passes over the nest, there were no statistically significant differences, however there was evidence for one outlier dam among the THC group who moved substantially (SFig 3b). Her pups were subsequently examined for evidence of outliers, but none were detected in later analyses. These data suggest that maternal behavior is not sufficiently altered to explain subsequent differences in offspring development. The model is as follows:

Maternal behavior model: Y = 𝝱_0_ + 𝝱_1_PND + 𝝱_2_conditionTHC+

𝝱_3_time-of-dayPM +𝝱_4_conditionTHC:PND +𝝱_5_conditionTHC:time-of-dayPM +𝝱_6_conditionTHC:PND:time-of-dayPM + **b**_1_subject + **ϵ**

Y= outcome measures (nest quality); 𝝱_i_= fixed effect coefficient; 𝝱_0_ = equation intercept; **b** =

random predictor; **ϵ** = random error; **:** = interaction;

conditionTHC= prenatal THC group relative to SAL as reference; PND = post-natal days (continuous); time-of-day = AM or PM

#### 5. Statistical Tests and Results

##### 5.1 *Embryo Body Volume*

To examine differences in body volume (mm^3^), an LMER was run with the interaction between treatment and sex as fixed effects and the litter size and coil as random effects. The results are below.

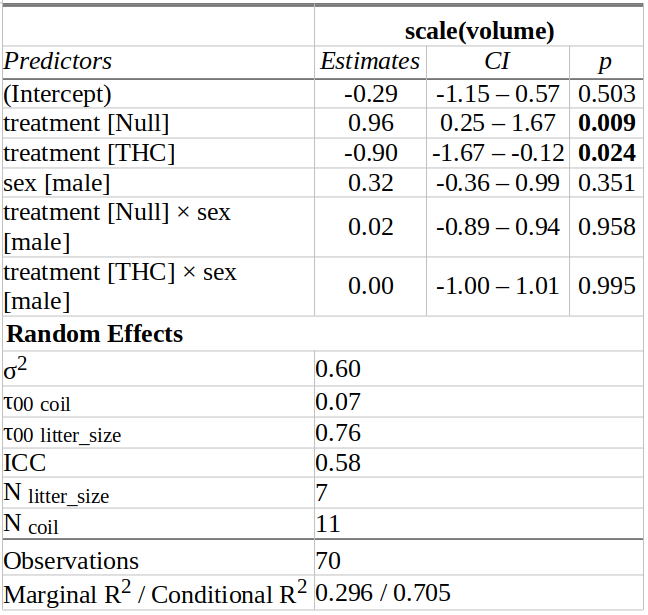

##### 5.2 *Neonate weights*

To examine the weight over time of pups, an LMER was run with the interaction between treatment, sex, and age as well as the main effect of whether the animals were scanned as fixed effects and the litter and subject ID as random effects. The results are below.

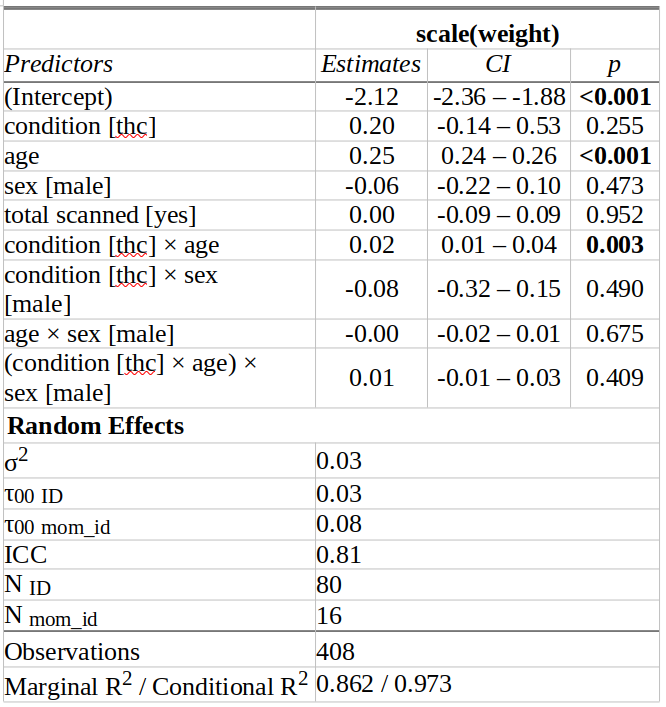

##### 5.3 *Adult weights*

To examine the weight over time of adolescent-adult offspring, an LMER was run with the interaction between treatment, sex, and age (modelled quadratically) as fixed effects and the litter and subject ID as random effects. The results are below.

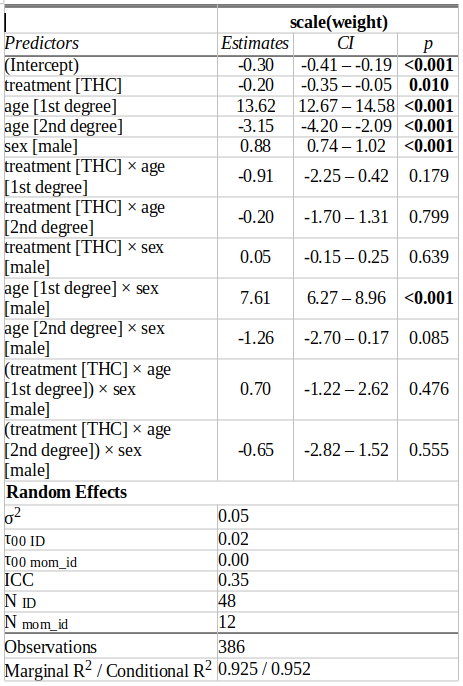

##### 5.4 Open Field Test

Total distance moved, time spent in the center, and passes through the center were examined with LMERs. Fixed effects were the interaction between treatment and sex, and the litter was the random effect. Significance was established with a Bonferroni corrected threshold of α = 0.05/3 = 0.016. The results are below.

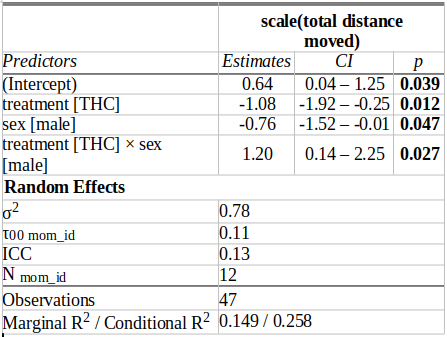

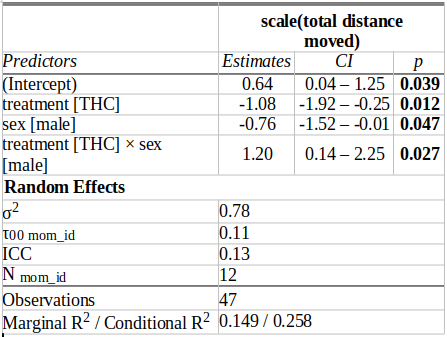

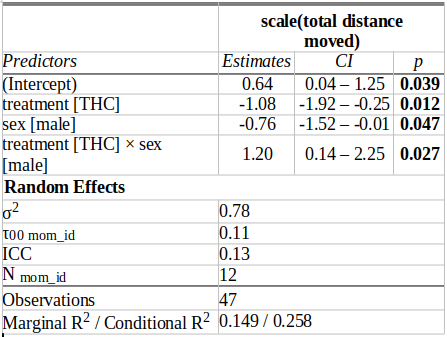

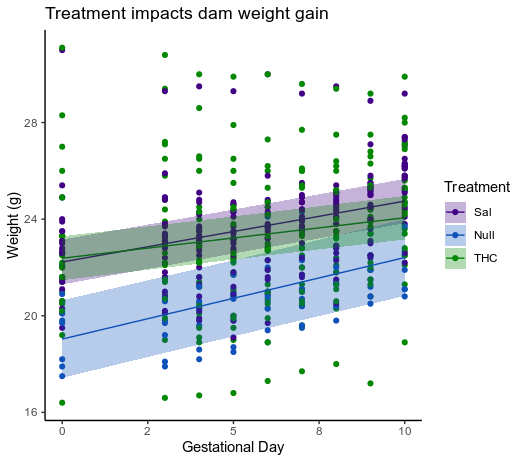

SFig 1. **Treatment impacts dam weight gain from GD 0 to 10.** Null dams (blue) gain weight more quickly, and start at a lower weight than Sal controls (purple). THC dams (green) gain weight less quickly than Sal controls.

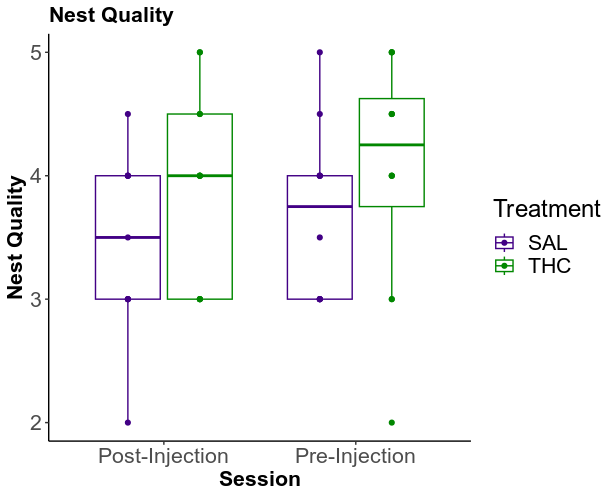

SFig 2. **Impact of PTE on quality of nests.** Nest quality, as assessed with an LMER, is unimpacted by either session (post-injection or pre-injection) or treatment.

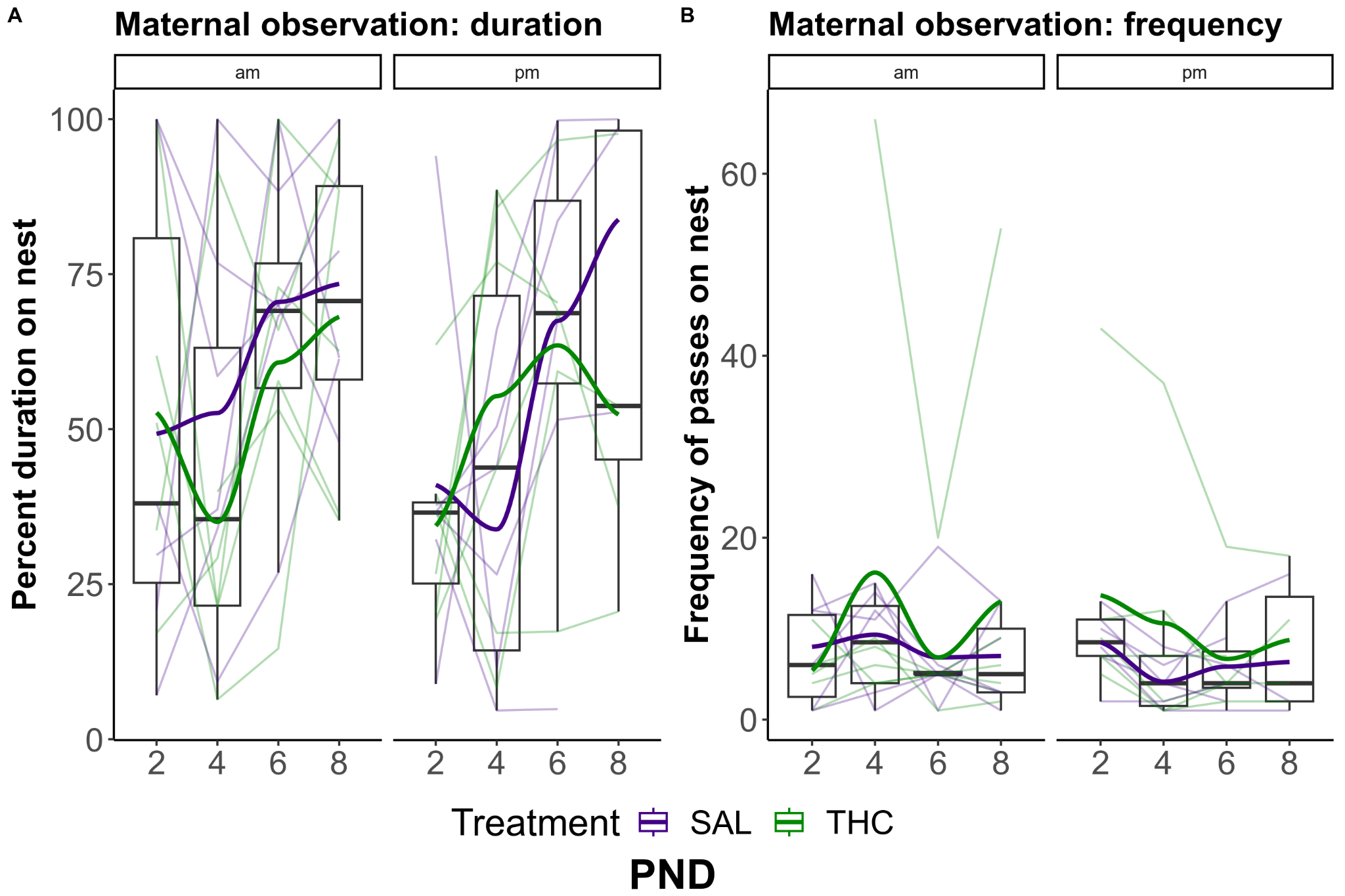

SFig 3. **Impact of PTE and PND on time spent on nest and duration of passes through nest.** Neither duration of time on nest (a), nor frequency of passes (b) differ by treatment. There is a trending effect of PND, such that dams spend more time on the nest later. Frequency of passes on nest reveals an outlier dam, who displayed hyperlocomotion.

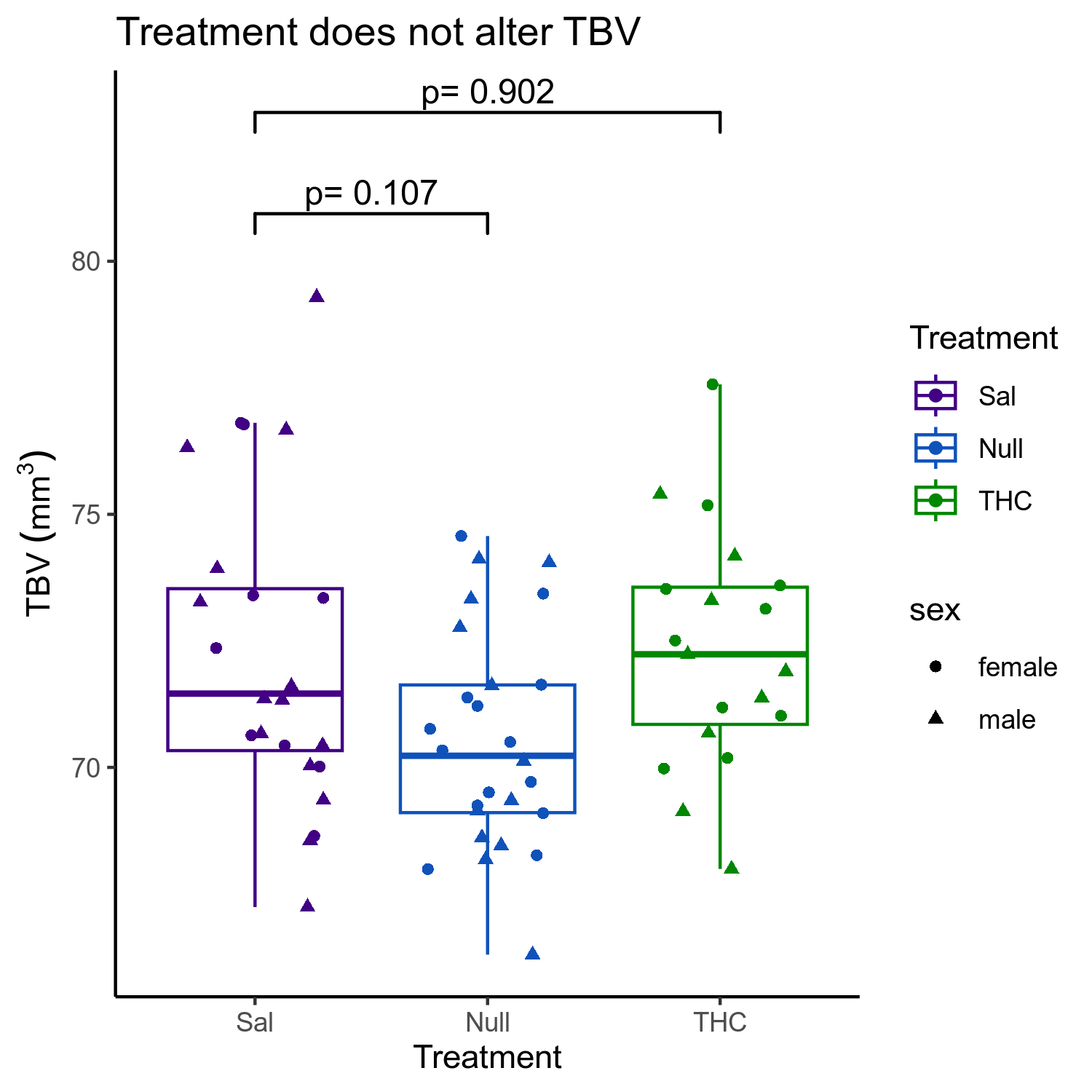

SFig. 4. **Impact of PTE on TBV of embryos.** TBV estimated from brain masks and compared with LMERs. No difference in TBV between treatments.

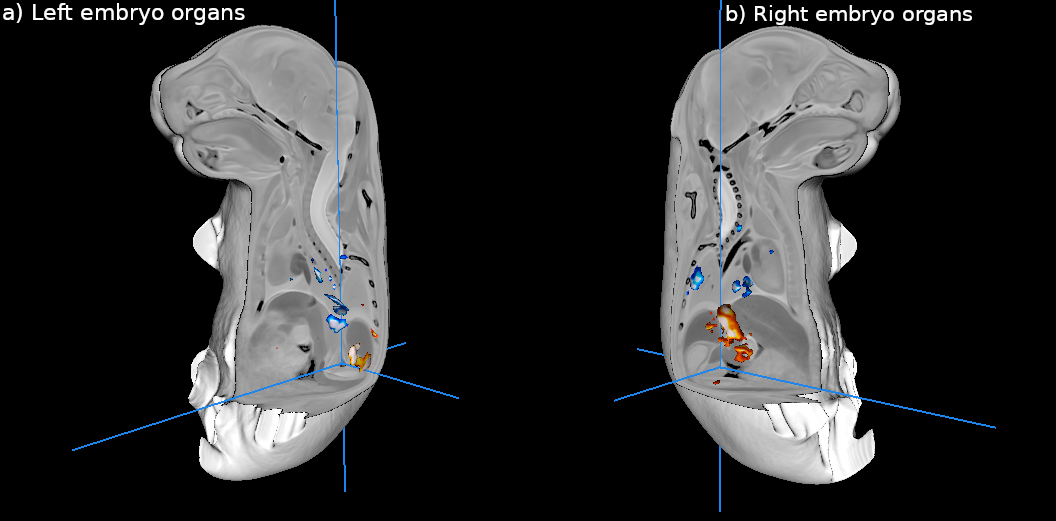

SFig. 5. **Impact of PTE on organ volume.** a) Embryo organ differences in THC compared to Sal groups, thresholded with FDR from 10%. b) The same map with a cut-out displaying the left side of the body. Color bars represent t-stats thresholded with FDR such that regions in blue indicate smaller volume in THC compared to Sal and warm colors indicate larger volume in THC compared to Sal. THC embryos showed larger volume in the liver and mixed results (clusters of larger and smaller voxels) in the lungs.

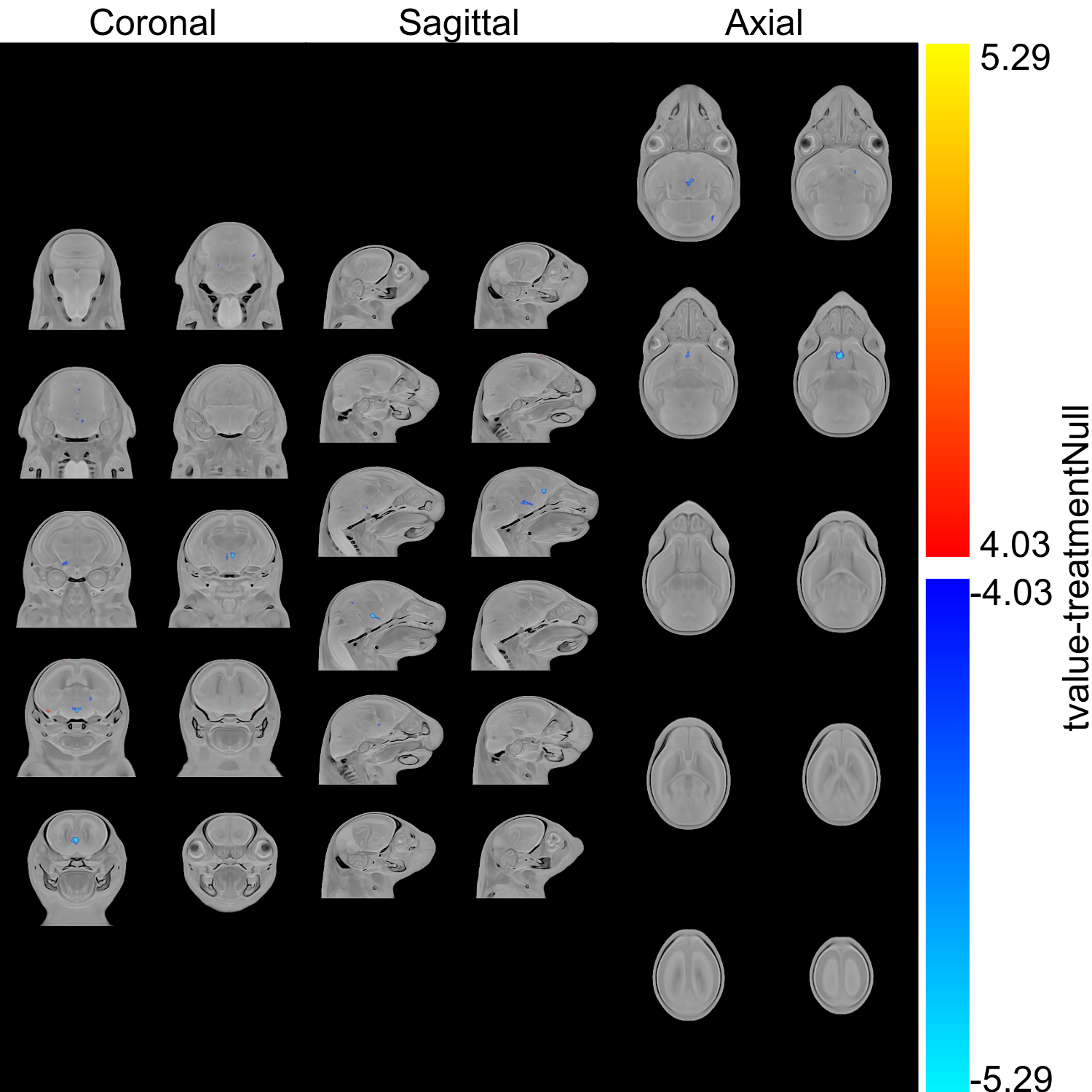

SFig. 6. **Minimal differences between Null and Sal embryo brains.** Heatmap showing differences between Null and Sal embryo brains, thresholded 5% FDR

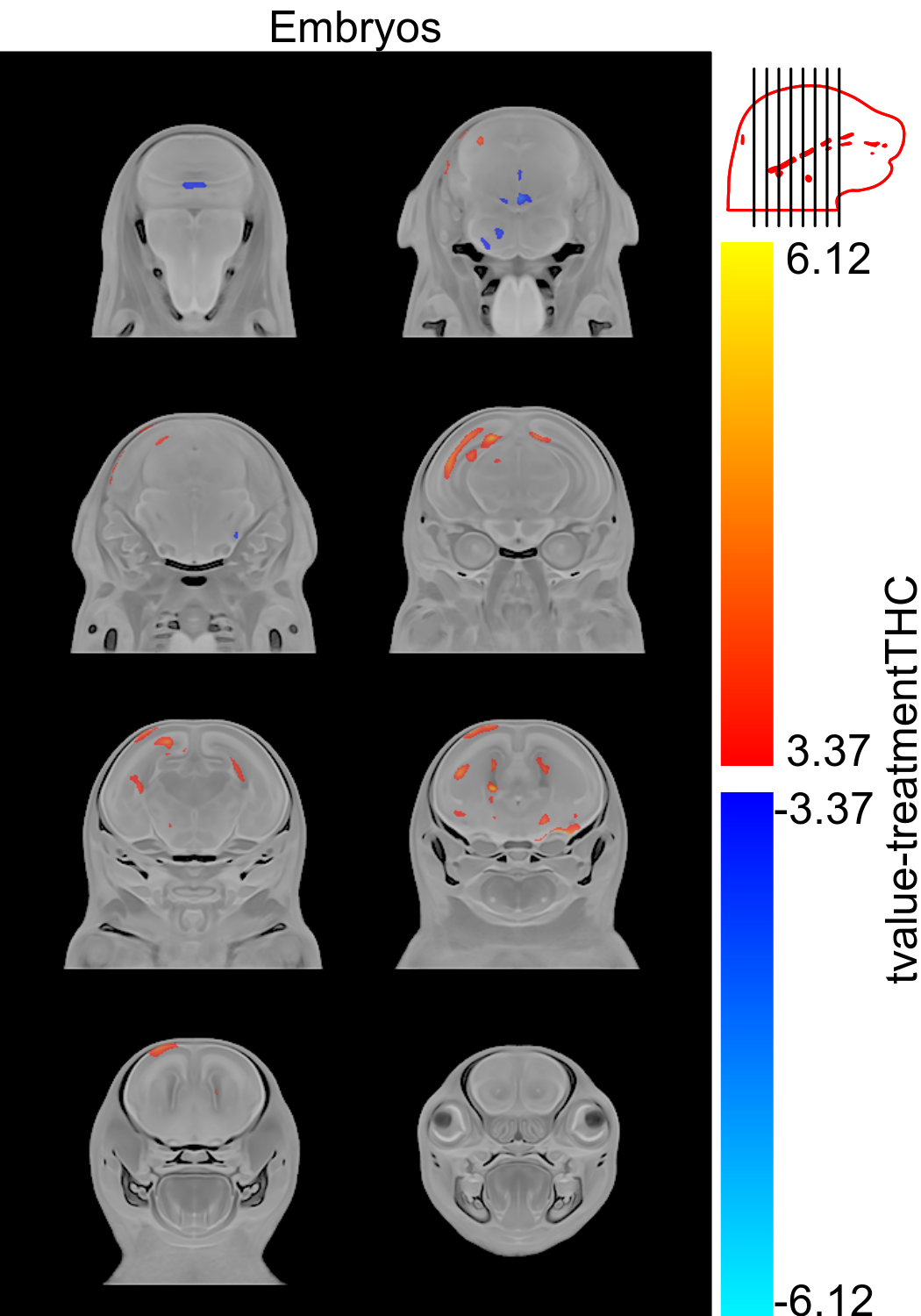

SFig. 7. **Impact of PTE on brain volume (absolute Jacobians).** While relative Jacobians are presented in the main text, here we present absolute Jacobians of embryos thresholded at 5% FDR with regions in warm colors showing larger volume in THC embryos and regions in blue showing smaller volume in THC embryos

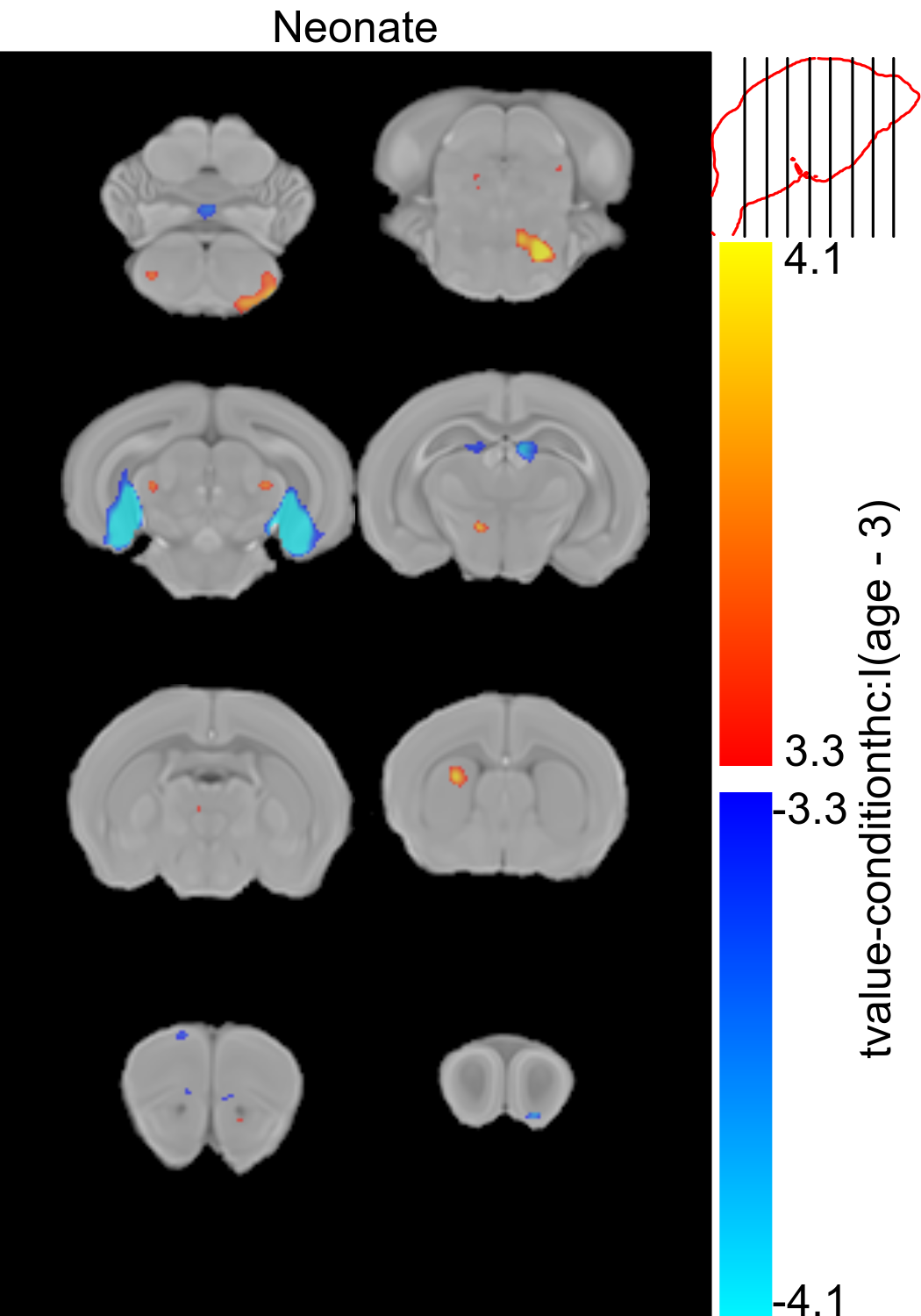

SFig. 8. **Impact of PTE on brain growth (relative Jacobians).** Relative Jacobians of neonates showing the age by condition interaction term thresholded at 5% FDR. Regions in warm colors show larger growth rate in THC pups and regions in cool colors show smaller growth rate in THC pups.

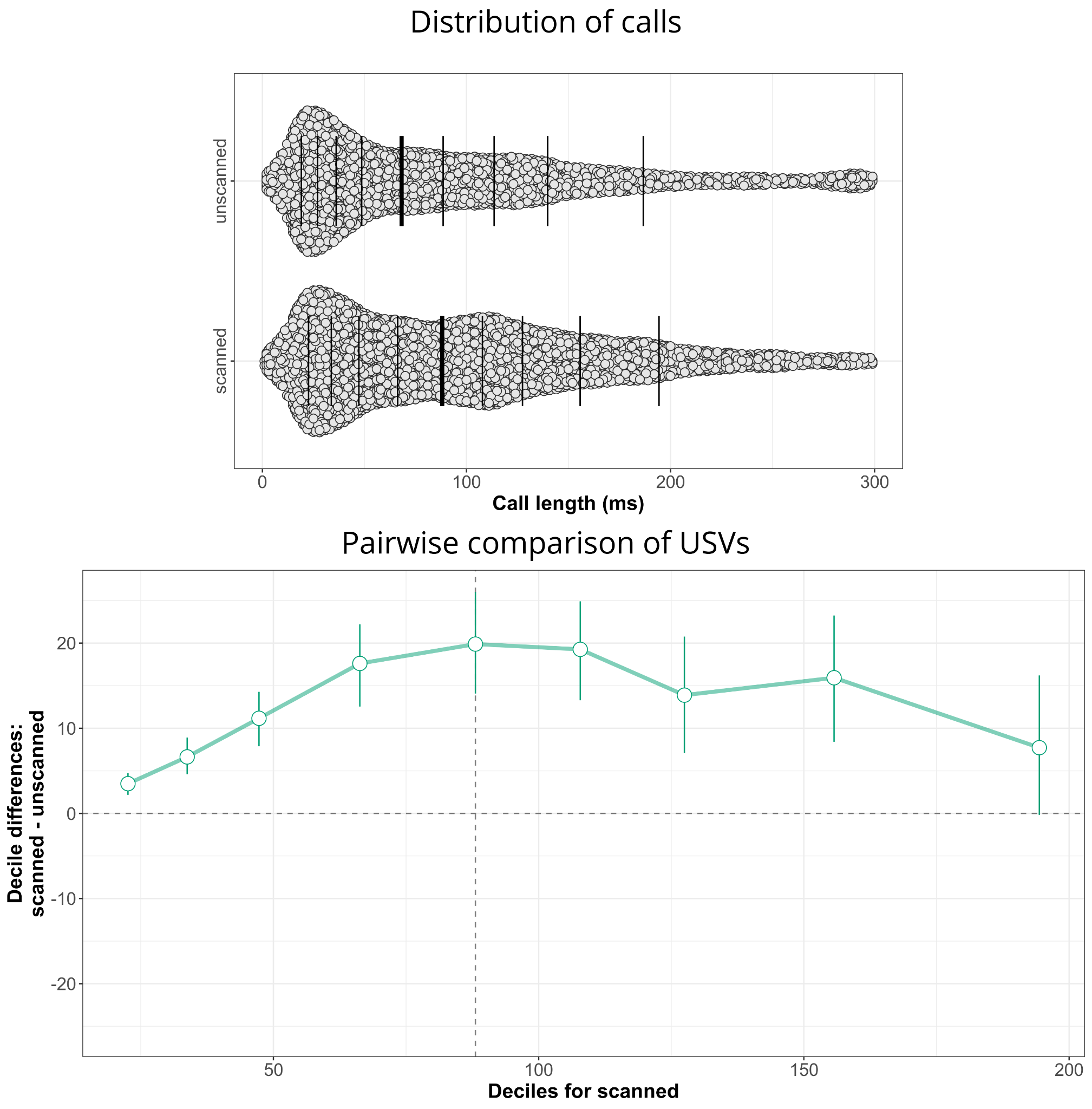

SFig 9. **Impact of scanning on USVs.** Top, distribution of calls between unscanned and scanned pups. Bottom, pairwise comparison between distributions for scanned and unscanned pups displaying increased calling in scanned pups.

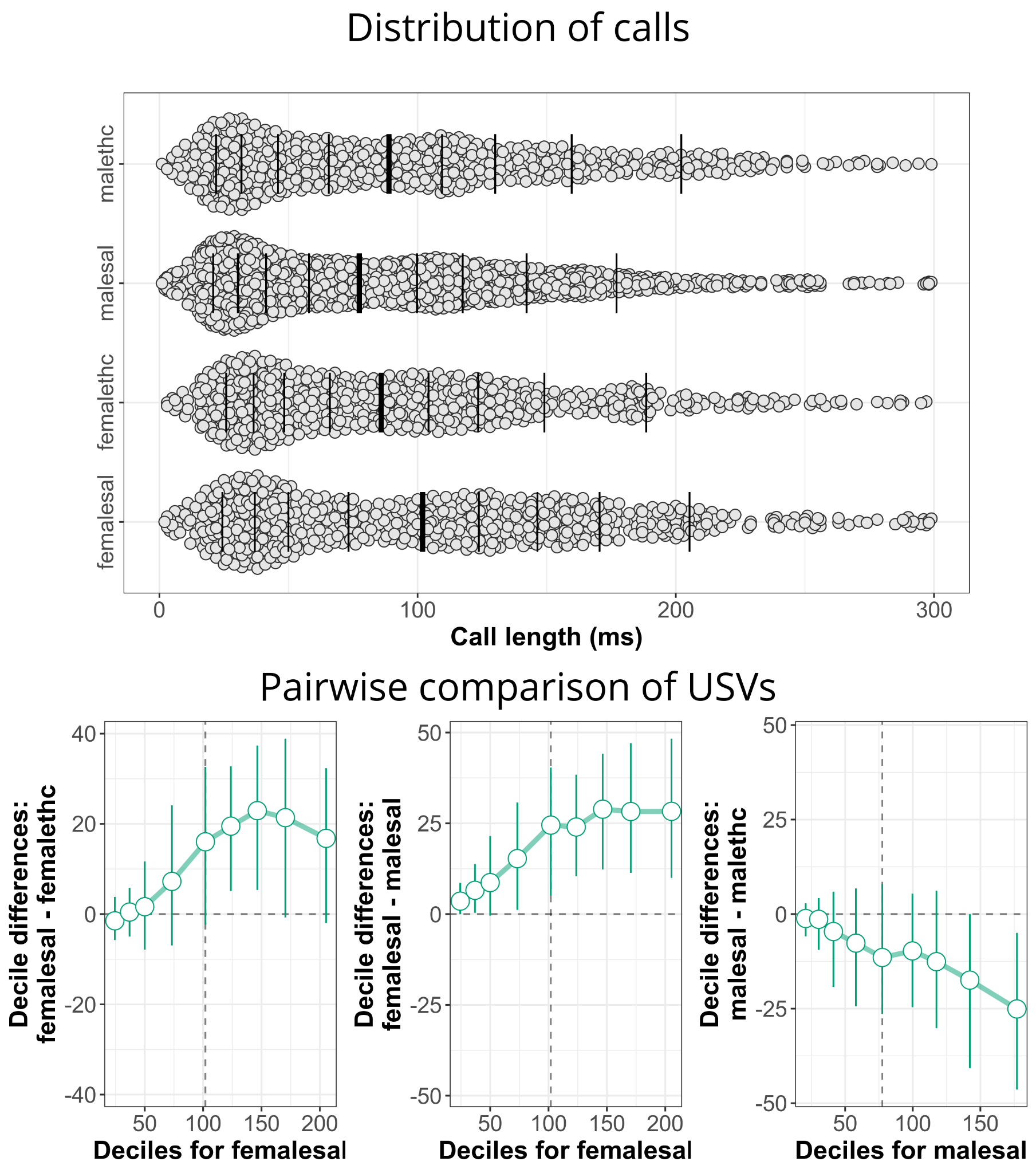

SFig 10. **Impact of PTE on USVs in MnCl2 subsample only.** Top: Distribution of calls divided by sex and condition for MnCl2 exposed pups only. Bottom, pairwise comparisons for female Sal vs female THC, female Sal vs male Sal, and male Sal vs male THC.

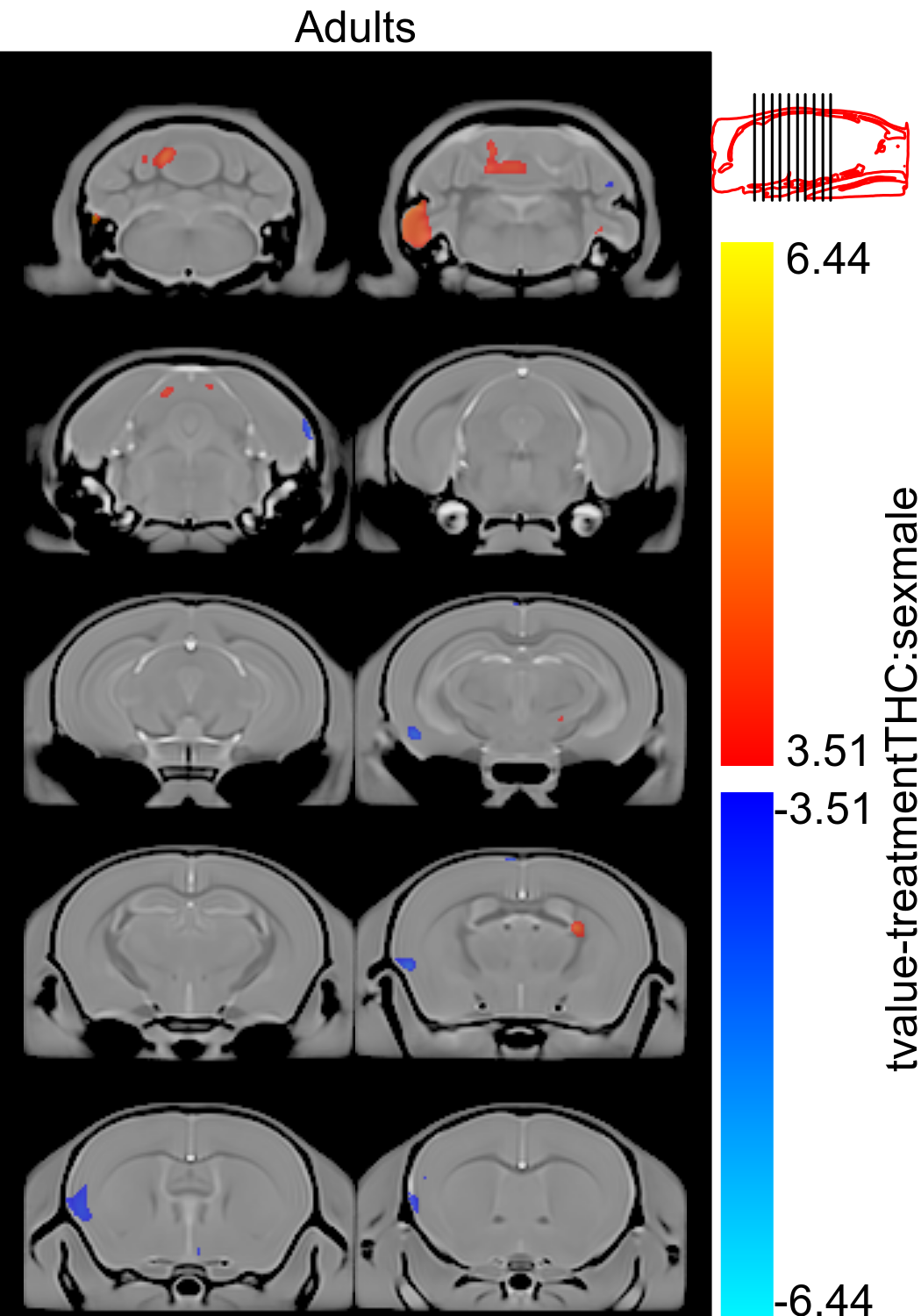

SFig. 11. **Impact of PTE and sex on brain volume (relative Jacobians).** Relative Jacobians from adults showing the treatment by sex interaction term thresholded at 5% with regions in warm colors showing larger volume in THC males and regions in blue showing smaller volume in THC males.

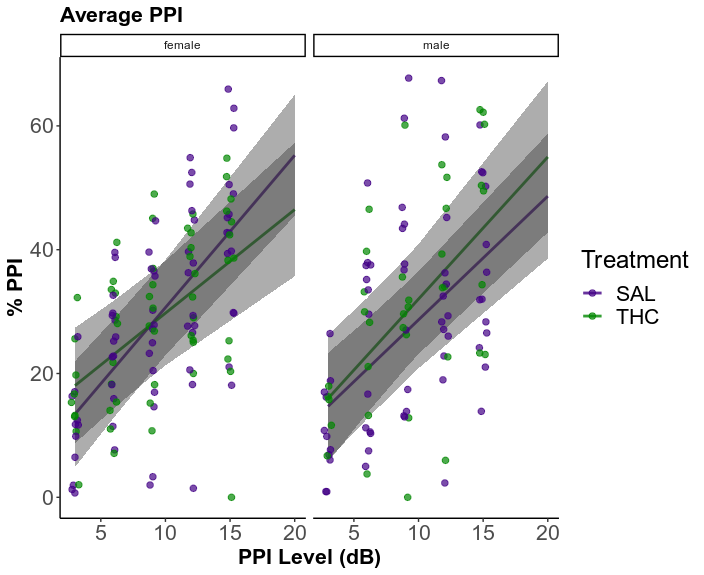

SFig 12. **Average %PPI by condition.** Trending reduction in %PPI for THC animals compared to Sal controls.
